# Free-Energy and Molecular Dynamics Analysis of Ser195 and Thr195 Rotameric States in Thrombin

**DOI:** 10.64898/2026.10.13.758922

**Authors:** Stephanie M. Sauve, Parth Chaturvedi, Mahmoud Moradi

## Abstract

Ser195 is highly conserved as the catalytic nucleophile in trypsin-like serine proteases, yet the molecular basis for the strong preference for serine over the chemically similar threonine residue remains unclear. Here, we used well-tempered metadynamics and microsecond-long molecular dynamics (MD) simulations to characterize the conformational energetics of S195 and T195 in thrombin. Well-tempered metadynamics revealed distinct rotameric free-energy landscapes in both the apo and the ligand-bound systems. In the apo state, S195 preferentially occupied a lower-angle rotameric state, while T195 exhibited two low-energy states with similar free energies. Unbiased simulations further showed greater conformational heterogeneity and increased higher-angle sampling for T195. Evaluation of the His57–residue 195 hydrogen bond further revealed that S195 maintained close hydrogen bond distances across both rotameric regions, whereas T195 exhibited longer and more variable His57–hydroxyl distances with reduced distance-based occupancy. Analysis of the local environment showed that the additional T195 methyl group was associated with redistributed protein contacts. Additional rotamer-conditioned analyses demonstrated differences in local packing within corresponding conformational states. Furthermore, solvent accessibility and water-contact analyses revealed that the effects of the substitution on the catalytic side-chain environment depend on rotameric conformation. Together, these findings demonstrate that two complementary simulation approaches converge on similar conformational preferences and suggest that the S195T substitution alters the rotameric free-energy landscape in association with changes in catalytic geometry, local packing, and solvation. These results provide insight into the evolutionary preference for serine over threonine at the catalytic position of thrombin.

## Introduction

Serine proteases are a widespread class of proteolytic enzymes that use a catalytic serine residue as a nucleophile to hydrolyze peptide bonds.^1,2^ Specifically, catalysis in trypsin-like serine proteases is mediated by a highly conserved catalytic triad consisting of Asp102, His57, and Ser195.^1,3–9^ In this charge relay system, Asp102 forms a hydrogen bond with His57 and helps to orient and to stabilize the histidine residue, enabling His57 to act as a general base during catalysis. His57 then promotes proton abstraction from the hydroxyl group of Ser195, facilitating nucleophilic attack on the substrate carbonyl carbon.^1,7,10^ The resulting tetrahedral intermediate is stabilized by the oxyanion hole formed by backbone atoms near the catalytic residue.^11,12^ Efficient proteolysis therefore depends on the precise organization and conformational dynamics of the catalytic triad.^4,13^

Despite the widespread conservation of Ser195 in trypsin-like proteases as the catalytic residue,^14^ the chemical basis for the strong evolutionary preference for serine over threonine at this position remains incomplete. Threonine possesses a hydroxyl group capable of acting as a nucleophile and is chemically similar to serine, differing by the presence of an additional methyl group on the residue’s side chain.^8^ Despite the strong similarities between the residues, naturally occurring trypsin-like proteases containing threonine in place of the catalytic serine have not been identified.^8,12^ Previous computational^15^ and observational studies^8,16^ proposed that the additional methyl group present in threonine introduces unfavorable steric interactions within the tightly packed active site which impair catalysis. However, this explanation alone does not establish how the methyl group alters the conformational energetics and local molecular environment of the catalytic side chain.

Structural and functional studies of the S195T mutant of thrombin provided important insight into this question. Pelc et al.^12^ determined high-resolution crystal structures of T195 thrombin in both the apo state and in complex with the irreversible inhibitor *H −D−Phe−Pro−Arg−CH*_2_*Cl* (PPACK). They found that the S195-to-T195 mutation reduced but did not abolish catalytic activity. These observations suggested that the S195T substitution may involve changes in the conformational preferences of the catalytic residue. Although crystal structures provide static structural snapshots,^17,18^ they do not directly reveal the free-energy landscape governing transitions among these conformations or how their populations evolve over time.^19^

Here, we use enhanced-sampling^20–22^ and microsecond-long unbiased MD simulations^23,24^ to characterize the conformational energetics of S195 and T195 in the thrombin protein. Well-tempered metadynamics was used to determine the side-chain rotameric free-energy landscapes of S195 and T195 in both apo and PPACK-bound thrombin, while independent simulations were performed for apo systems to examine conformational sampling over extended timescales. We examined the hydrogen bond between residue 195 and His57 to investigate how the S195T substitution influences catalytic-triad geometry. Additionally, we further quantified local protein contacts, minimum distances, solvent accessibility, and water interactions surrounding residue 195 to determine how the additional methyl group of T195 modifies the local molecular environment compared to S195. Together, these analyses provide a dynamic and energetic framework for understanding how the S195-to-T195 substitution reshapes the catalytic side-chain ensemble and why serine is preferentially selected at the nucleophilic position of the thrombin active site.

## Methods

Residue numbering throughout the manuscript follows conventional chymotrypsinogen numbering.^25^

### System Preparation

We designed four thrombin simulation systems based on high-resolution crystal structures representing the mutant T195 apo state (PDB entry: 4RKJ, 1.70 Å resolution)^12^ and the mutant T195 PPACK-bound state (PDB entry: 4RKO, 1.84 Å resolution).^12^ Representations of the created systems are shown in Figure 1 which was generated using Visual Molecular Dynamics (VMD).^26^ For the apo S195 and the PPACK-bound S195 systems, residue 195 was mutated back to the wild-type amino acid residue. For all systems, we utilized a Monte Carlo algorithm to model a missing loop region using the program MODELLER,^27^ with crystal structures modeled between residues 148-149E. The resulting topology and coordinates were used to generate the apo S195 (Fig. 1A) and T195 systems (Fig. 1B). The prepared apo thrombin systems were made using the Solution Builder module in CHARMM-GUI^28^ in which 0.15 M NaCl was included (in addition to the counterions used to neutralize the protein) to mimic physiological conditions. The simulation systems were solvated in a cubic box of TIP3P water molecules^29^ (apo thrombin S195 systems had dimensions of 86 Å *×* 86 Å *×* 86 Å and apo thrombin T195 systems had dimensions of 86 Å *×* 86 Å *×* 86 Å). The total number of atoms in the apo thrombin systems were 59,474 atoms for S195 and 59,522 atoms for T195.

**Fig. 1.**
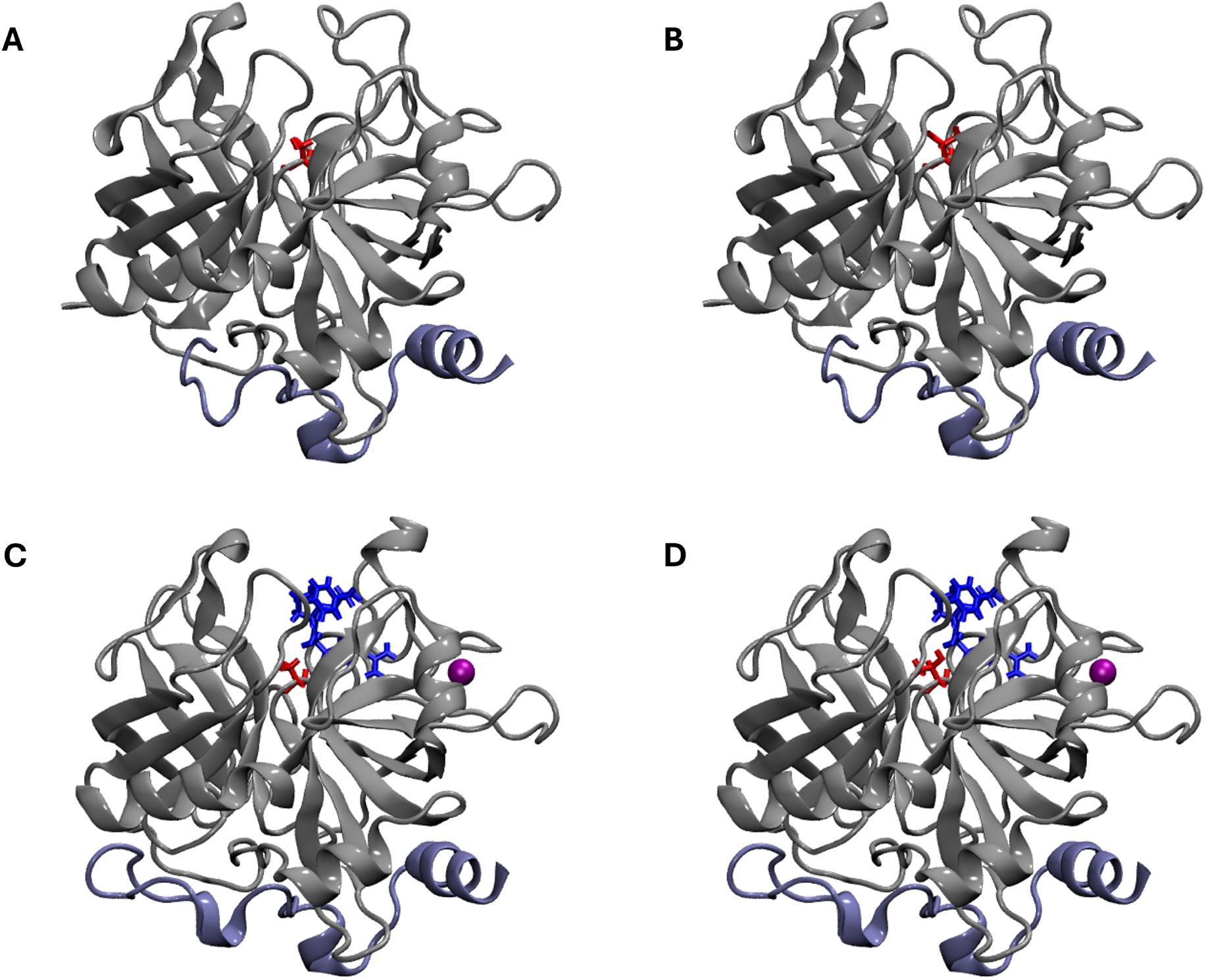
Representative structures of the four simulated systems are shown: (A) apo S195, (B) apo T195, (C) PPACK-bound S195, and (D) PPACK-bound T195. The thrombin heavy chain is shown in a gray NewCartoon representation, and the light chain is shown in an ice-blue NewCartoon representation. The mutated residue at position 195 is shown in red bonds representation. For the ligand-bound systems, PPACK is shown in blue bonds representation and the sodium ion is shown in purple van der Waals (VDW) representation.

PPACK-bound thrombin systems were constructed using PSFgen.^26^ Standard N-terminus and C-terminus patches were applied to the PPACK peptide during system construction. Since PPACK contains a D-amino-acid residue, the CHARMM36m D-amino-acid stream file, containing the required topology and parameter definitions, was included during system construction. Additionally, a custom topology patch was subsequently applied to reproduce the covalent linkage between PPACK and thrombin at His57 and the C-terminal Arg residue of PPACK. The thrombin-associated Na^+^ ion was also retained in the constructed systems. The resulting topology and coordinates were used to generate the PPACK-bound S195 (Fig. 1C) and T195 systems (Fig. 1D).

The prepared PPACK-bound thrombin systems were made using PSFgen in which 0.15 M NaCl was included (in addition to the counterions used to neutralize the protein) to mimic physiological conditions. The simulation systems were solvated in a cubic box of TIP3P water molecules^29^ (PPACK-bound thrombin S195 systems had dimensions of 100 Å *×* 100 Å *×* 100 Å and PPACK-bound thrombin T195 systems had dimensions of 100 Å *×* 100 Å *×* 100 Å). The total number of atoms in the PPACK-bound thrombin systems were 94,990 atoms for S195 and 94,996 atoms for T195.

### Well-Tempered Metadynamics Simulations

Prior to running well-tempered metadynamics,^20–22^ each system was energy-minimized, equilibrated, and subjected to an unbiased production simulation. All MD simulations were performed in NAMD 2.14^30,31^ using the CHARMM36m force field.^32^ For the PPACK-bound systems, the thrombin-associated Na^+^ ion was maintained within its coordination site using two harmonic distance restraints between Na^+^ and coordinating oxygen atoms of Arg221A and Lys224.^33–37^ The restraints were implemented using the NAMD extraBonds^30^ feature with a force constant of 10.0 kcal mol^−1^ Å^−2^ and equilibrium distances of 2.25 and 2.23 Å, respectively. Energy minimization was initially applied to the thrombin containing systems using the conjugate gradient algorithm for 10,000 steps. The systems were subsequently equilibrated for 15 ns in the NVT ensemble, followed by a 100 ns unbiased production simulation in the NPT ensemble. A time step of 2 fs was utilized throughout the simulations. The temperature was maintained at 310 K using a Langevin thermostat and the Nośe-Hoover Langevin piston method^38^ was used to control pressure at 1 bar. One independent production trajectory was generated for each of the four systems, and the final configurations from these simulations were used as the starting structures for the subsequent well-tempered metadynamics simulations.

Well-tempered metadynamics simulations were subsequently performed using the collective variables (Colvars) module implemented in NAMD^39^ for 100 ns for each of the four thrombin systems. The simulations were conducted in the NPT ensemble at 310 K and 1 bar using the same thermostat and barostat settings described above. The C–C*α*–C*β*–O*γ* dihedral angle of residue 195 was selected as the collective variable to characterize the free-energy landscape of the catalytic side chain.^12^ The dihedral angle was defined over the range of *−*180*^◦^* to 180*^◦^* with a collective-variable width of 15*^◦^*. Gaussian hills with an initial height of 0.2 kcal/mol and a width of 15*^◦^* were deposited every 500 simulation steps, corresponding to 1 ps. Well-tempered metadynamics was performed using a bias temperature of 1550 K for a biasing factor of 6. Collective-variable trajectories were written every 500 simulation steps.

The convergence of the calculated free-energy profiles was evaluated by comparing the potential of mean forces (PMFs) obtained throughout the final 20 ns of each well-tempered metadynamics simulation (Figure **??**) and by evaluating the temporal stability of selected regions of the free-energy landscape (Figure **??**). Free-energy profiles from the final 20 ns of each 100 ns well-tempered metadynamics simulation were used to obtain the reported PMFs. Twenty PMF profiles spanning this interval were analyzed. Prior to averaging, each individual PMF was referenced to its respective free-energy minimum by setting the minimum to 0 kcal/mol. The dihedral-angle coordinate was converted from the *−*180*^◦^*-180*^◦^* representation to 0*^◦^*-360*^◦^* and the PMFs were averaged at each dihedral-angle value. The sample standard deviation across the 20 PMF profiles was calculated at each angle and is shown as the shaded region in the free-energy plots (Fig. 2). The resulting mean PMF was subsequently shifted such that its minimum free energy was 0 kcal/mol.

**Fig. 2.**
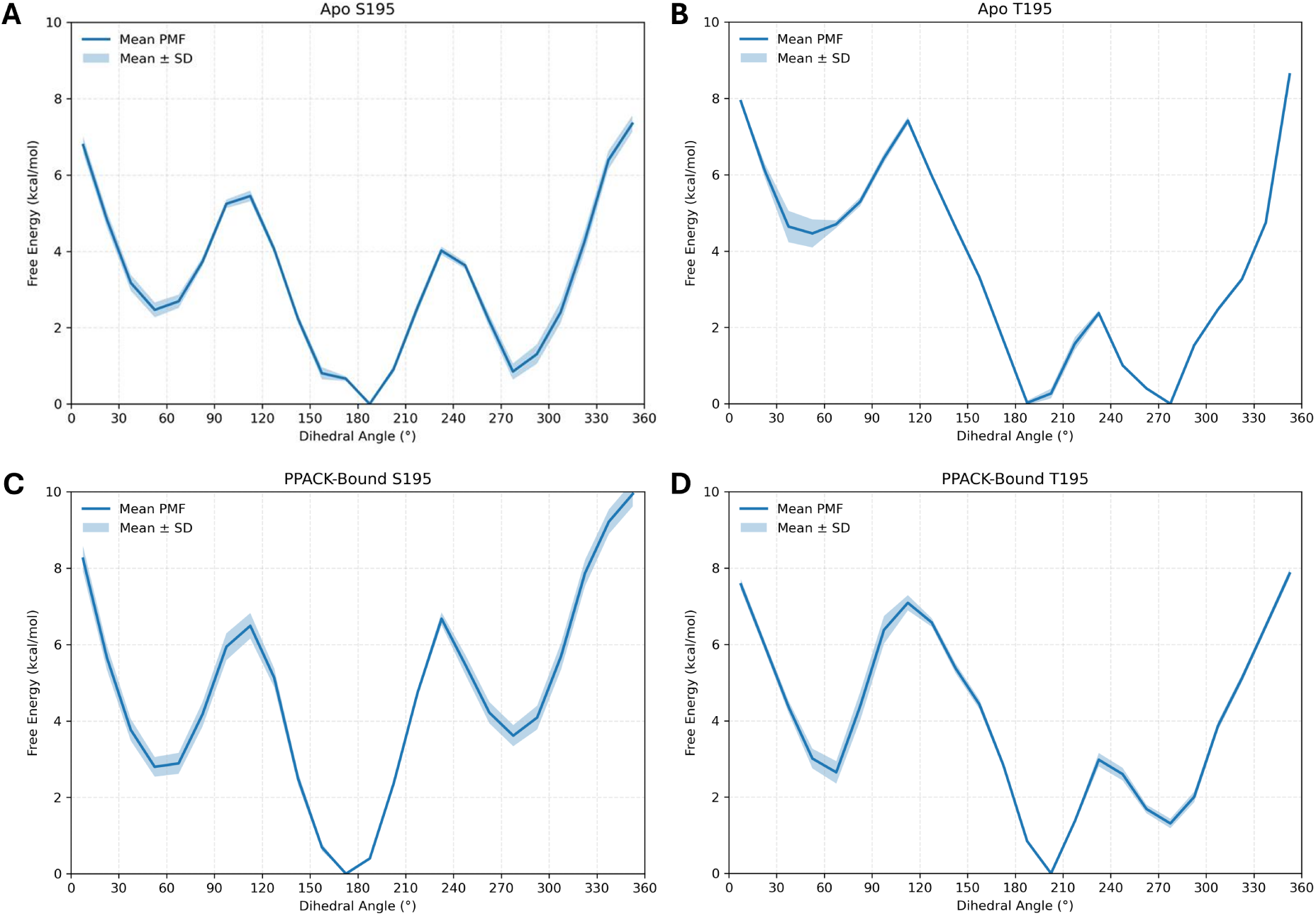
Free-energy profiles of the S195 and T195 side-chain rotameric states in apo and PPACK-bound thrombin. PMFs were obtained from 100 ns well-tempered metadynamics simulations using the C–C*α*–C*β*–O*γ* dihedral angle of residue 195 as the collective variable. The PMFs shown represent the average over the final 20 ns of each simulation, with shaded regions indicating the standard deviation over this averaging interval. Profiles are shown for (A) apo S195, (B) apo T195, (C) PPACK-bound S195, and (D) PPACK-bound T195. Free energies were referenced to the minimum of each individual PMF.

### Long-Timescale MD Simulations

Prior to the microsecond-long simulations, the apo S195 and apo T195 systems prepared using CHARMM-GUI were first energy-minimized for 10,000 steps using the conjugate-gradient algorithm. The systems were then equilibrated for 15 ns, followed by a 15 ns unbiased production simulation on the Frontera supercomputer. The same general simulation conditions described above were used, including NAMD 2.14, the CHARMM36m force field, a temperature of 310 K, and a pressure of 1 bar during the NPT stage. The final configurations from these 15 ns production simulations were subsequently used to initiate the long-timescale simulations.

Five independent long-timescale simulations were performed in the NPT ensemble for each apo thrombin system using Anton 3^40^ with the Desmond MD framework. For both S195 and T195, five independent replicates were simulated for 30 *µ*s each, corresponding to an aggregate sampling time of 150 *µ*s per system and 300 *µ*s across both systems. Simulations were performed using a 2.5 fs integration time step at a temperature of 310 K and a reference pressure of 1 bar. Temperature was controlled using the Antithetic thermostat^41^ and pressure was maintained using a Monte Carlo barostat.^42^

### Trajectory Analysis

#### Dihedral-Angle and Free-Energy Analysis

The conformational sampling of residue 195 in the long-timescale apo simulations was characterized using the C–C*α*–C*β*–O*γ* side-chain dihedral angle.^12^ Dihedral angles were represented over the range of 0-360*^◦^* and were calculated for each of the five independent 30 *µ*s trajectories for both the S195 and T195 systems. For visualization of the replicate-level distributions, normalized histograms were constructed using 5*^◦^* bins and smoothed using a Gaussian filter with a standard deviation of 1.5 bins while treating the angular coordinate as periodic. The five trajectories for each system were also pooled to obtain combined distributions corresponding to an aggregate sampling time of 150 *µ*s per system.

Population-derived free-energy profiles were calculated from the pooled dihedral-angle distributions of the five replicates. The 0-360*^◦^* dihedral-angle range was divided into 36 bins of 10*^◦^*, and the probability, *P* (*θ*), of each bin was calculated from its normalized population. Free energies were then calculated according to the Boltzmann relation:^43^

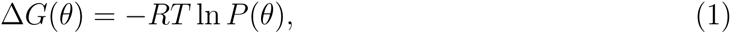

where *R* = 1.9872 *×* 10*^−^*^3^ kcal mol*^−^*^1^ K*^−^*^1^ and *T* = 310 K. Bins with zero observed population were masked and excluded from the free-energy profiles to avoid undefined values arising from ln(0). Each resulting free-energy profile was shifted such that its minimum was set to 0 kcal/mol. Free-energy profiles were displayed over the range of 0–10 kcal/mol.

For quantitative comparison of the metadynamics and unbiased MD free-energy profiles, the lower- and higher-angle conformational basins were identified within the angular ranges of 150–210*^◦^* and 260–310*^◦^*, respectively. The position and free energy of each basin were determined from the local minimum within its corresponding angular range. The relative free-energy difference between the basins was calculated by subtracting the free energy of the lower-angle minimum from that of the higher-angle minimum. Forward and reverse barrier heights were calculated relative to the corresponding basin minima using the maximum free energy in the intervening region of 210–260*^◦^*.

#### Hydrogen Bond Analysis

Hydrogen bond analysis was conducted using the VMD HBond plugin,^26^ with a cutoff distance of 4.0 Å and an angle cutoff of 30*^◦^*. Specifically, to characterize the positioning of the catalytic hydroxyl relative to His57, the distance between His57 N*ɛ*2 and the hydroxyl oxygen of residue 195 (S195 OG or T195 OG1) was analyzed. The minimum distance of the atoms involved in the hydrogen bond was examined over time for each replicate. To assess its dependence on side-chain conformation, distance measurements were matched to the corresponding dihedral angles using trajectory frame indices and analyzed separately within the lower-angle (150-210*^◦^*) and higher-angle (260-310*^◦^*) rotameric regions. For each replicate and rotameric region, the median His57-O195 distance was calculated, along with the percentage of frames in which the distance was within 4 Å. Frames outside these regions were excluded from the rotamer-conditioned analysis. The replicate-level median distances and distance-based contact occupancies were summarized as the mean and standard deviation across five independent simulations.

#### Residue Contact and Minimum-Distance Analysis

To characterize changes in the local protein environment surrounding residue 195, residue-level contacts were calculated for the S195 hydroxyl oxygen (OG), the T195 hydroxyl oxygen (OG1), and the additional T195 methyl carbon (CG2). A contact was defined when at least one atom of a neighboring protein residue was within 5 Å of the specified residue-195 atom. Residue 195 itself was excluded from the contact selection. Contact occupancy was calculated as the percentage of trajectory frames in which a given neighboring residue formed at least one contact with the target atom. Contact occupancies were calculated independently for each of the five replicates, and replicate means and standard deviations were subsequently determined.

Minimum-distance analyses were performed for the neighboring residues S214, V213, and G216, which exhibited prominent differences in contact occupancy between the S195 and T195 systems. For each trajectory frame, the minimum distance between S195 OG, T195 OG1, or T195 CG2 and any heavy atom of the selected neighboring residue was calculated. Each trajectory was summarized by its median minimum distance. The five replicate-level median values were then used to calculate the mean and standard deviation for each residue-195 atom and neighboring-residue pair.

To determine whether differences in local interactions between S195 and T195 were associated with changes in rotameric populations, contact occupancies and minimum distances were additionally analyzed separately for the lower-angle (150-210*^◦^*) and higher-angle (260-310*^◦^*) rotameric regions. Trajectory frames were classified according to the C–C*α*–C*β*–O*γ* dihedral angle of residue 195, and the corresponding minimum-distance measurements were assigned to each rotameric region using matching frame indices. Frames outside these angular ranges were excluded from the rotamer-conditioned analyses. For each replicate and rotameric region, contact occupancy was calculated as the percentage of frames with a minimum heavy-atom distance below 5 Å, while the median minimum distance was calculated from all frames within that region. The mean and standard deviation of the resulting replicate-level measurements were then calculated across the five independent simulations.

#### Solvent Accessibility and Water-Contact Analysis

Atom-specific solvent-accessible surface area (SASA)^44^ was calculated for S195 OG, T195 OG1, and T195 CG2 using a solvent probe radius of 1.4 Å. For each of the five independent trajectories for each system, the median atom-specific SASA was calculated, and the replicate-level median values were subsequently used to determine the mean and standard deviation for each atom.

Local water contact was quantified by determining the presence of water oxygen atoms within 3.5 Å of S195 OG, T195 OG1, or T195 CG2. Water-contact occupancy was defined as the percentage of trajectory frames containing at least one water oxygen within this cutoff. Occupancies were calculated independently for each replicate, and the mean and standard deviation were calculated across the five replicates.

To assess the dependence of solvent accessibility and hydration on side-chain conformation, atom-specific SASA and water-contact occupancies were additionally calculated separately for the lower-angle (150-210*^◦^*) and higher-angle (260-310*^◦^*) rotameric regions. For each replicate, SASA and water-contact measurements were matched to the corresponding dihedral angles using trajectory frame indices. Frames outside the defined rotameric regions were excluded from these analyses. Within each rotameric region, the median atom-specific SASA was calculated for each replicate, while water-contact occupancy was determined as the percentage of frames containing at least one water oxygen within 3.5 Å of the corresponding atom. The resulting replicate-level median SASA values and water-contact occupancies were summarized as the mean and standard deviation across the five independent simulations.

## Results and Discussion

### Metadynamics Reveals Distinct Free-Energy Landscapes for S195 and T195

To characterize the free-energy landscape of the catalytic residue, well-tempered metadynamics simulations were performed using the C–C*α*–C*β*–O*γ* dihedral angle of S195 or T195 as the collective variable. Simulations were carried out for both the apo and PPACK-bound states, and the PMFs were averaged over the final 20 ns of each 100 ns simulation (Fig. 2).

In apo S195 thrombin, the global free-energy minimum was located at 187.5*^◦^*, with a second low-energy minimum at 277.5*^◦^* that was 0.85 kcal/mol higher in free energy (Fig. 2A). The two basins were separated by a free-energy maximum of approximately 4.0 kcal/mol near 232.5*^◦^*. Overall, the location of the global free-energy minimum for S195 is consistent with the findings reported by Pelc et al. in which the mean S195 dihedral angle was located near 156*^◦^* with a standard deviation of 37*^◦^*.^12^ In contrast, the apo T195 landscape exhibited minima at both 187.5*^◦^* and 277.5*^◦^*with nearly identical free energies (Fig. 2B). The 187.5*^◦^* state was only 0.02 kcal/mol above the minimum at 277.5*^◦^*, a difference smaller than the uncertainty of the calculated PMF. Moreover, the free-energy maximum separating these states was reduced to approximately 2.4 kcal/mol. Our findings regarding the location of a higher-angle free-energy minimum for T195 are similar to that of Pelc et al. in which the T195 dihedral angle was found to be near 305*^◦^*.^12^ Thus, substitution of S195 with T195 altered both the relative stability and energetic separation of the major rotameric states, converting the preferential 187.5*^◦^*basin observed for S195 into two low-energy states.

PPACK binding substantially redistributed the S195 free-energy landscape (Fig. 2C). The global minimum was located at 172.5*^◦^*, while the alternative minimum at 277.5*^◦^* increased to 3.61 kcal/mol above the global minimum, compared with only 0.85 kcal/mol in the apo state. The free-energy maximum between the two principal basins also increased to approximately 6.67 kcal/mol near 232.5*^◦^*. These results indicate that PPACK binding strongly biases S195 toward the lower-angle rotameric state.

PPACK binding also altered the T195 landscape, although the energetic discrimination between the two major rotameric states was less pronounced than for S195 (Fig. 2D). The global minimum occurred at 202.5*^◦^*, with the neighboring 187.5*^◦^* configuration remaining within 0.84 kcal/mol of the minimum. A secondary minimum persisted at 277.5*^◦^*, 1.31 kcal/mol above the global minimum, with an intervening free-energy maximum of approximately 2.98 kcal/mol near 232.5*^◦^*. Thus, PPACK binding shifted the relative stability of the T195 rotameric states toward the lower-angle basin, but the higher-angle state remained more energetically accessible than in PPACK-bound S195. Our observation that PPACK binding causes a shift towards lower-angle conformational states is consistent with the findings reported by Pelc et al. in which the binding of PPACK shifted the dihedral angle curve to a mean of 148*^◦^* with a standard deviation of 36*^◦^*.^12^

### Long-Timescale Apo Simulations Reveal Distinct S195 and T195 Rotameric Ensembles

To determine whether the differences observed in the well-tempered metadynamics free-energy landscapes were reflected in long-timescale conformational sampling of the apo enzyme, five independent 30 *µ*s unbiased MD simulations were analyzed for both S195 and T195 thrombin. The C–C*α*–C*β*–O*γ* dihedral angle was monitored throughout each trajectory, providing an aggregate sampling time of 150 *µ*s for each system.

The S195 simulations exhibited relatively consistent dihedral-angle distributions across the five independent replicates (Fig. 3A). Sampling was dominated by a lower-angle population centered approximately within the 150-180*^◦^* region with a less populated higher-angle state observed near 280-300*^◦^*. Overall, the lower-angle state remained the predominant feature of the S195 ensemble, again demonstrating consistency with the findings reported by Pelc et al. who reported the S195 dihedral angle to be near 148*^◦^*.^12^ In contrast, T195 displayed substantially greater replicate-dependent heterogeneity (Fig. 3B). Most of the T195 trajectories preferentially sampled the higher-angle region near 280-300*^◦^*, but one replicate predominantly sampled the lower-angle region near 180*^◦^*. These differences indicate that the T195 side chain can access both conformational regions but that their relative populations vary considerably among independent trajectories. Our findings that T195 samples the higher-angle region near 280-300*^◦^* is again consistent with the findings of Pelc et al. who showed the T195 dihedral angle to be near 305*^◦^*.^12^ Individual trajectory time series plots (Fig. **??**) further demonstrate that the broader T195 conformational ensemble arises from increased replicate-to-replicate variability and sampling of distinct dihedral states across the long-timescale simulations.

**Fig. 3.**
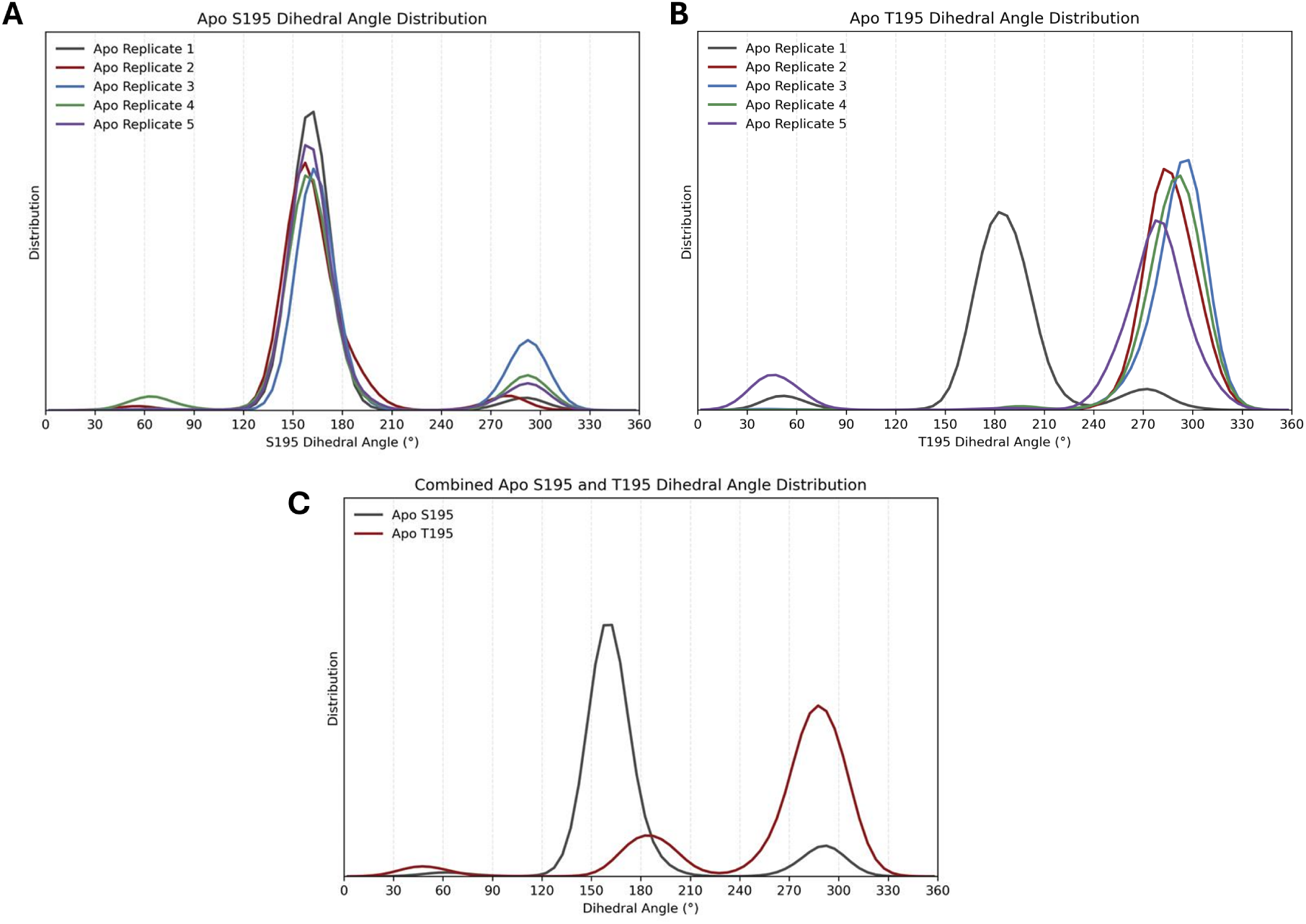
Dihedral-angle distributions of S195 and T195 during long-timescale apo thrombin simulations. Normalized distributions of the C–C*α*–C*β*–O*γ* dihedral angle are shown for five independent 30 *µ*s unbiased molecular dynamics simulations of (A) apo S195 and (B) apo T195 thrombin. Each curve represents an individual simulation replicate. Replicate 1 is shown in gray, replicate 2 is shown in red, replicate 3 is shown in blue, replicate 4 is shown in green, and replicate 5 is shown in purple. (C) Combined distributions obtained by pooling the five replicates for each system, corresponding to an aggregate sampling time of 150 *µ*s per system. S195 data is shown in gray while T195 data is shown in red. Dihedral angles were represented from 0–360*^◦^*, and distributions were calculated using 5*^◦^* bins followed by Gaussian smoothing.

Pooling the five trajectories for each system further highlighted the difference between the S195 and T195 ensembles (Fig. 3C). The combined S195 distribution was dominated by the lower-angle population with a smaller population at higher dihedral angles. In contrast, the combined T195 distribution exhibited a dominant higher-angle population together with a distinct lower-angle population. Thus, the S195-to-T195 substitution substantially altered the relative populations of the major conformational states sampled by the catalytic side chain in the apo enzyme.

The corresponding population-derived free-energy profiles further illustrated these differences (Fig. 4). The S195 profile contained a pronounced low-free-energy basin in the lower-angle region and a secondary minimum near the higher-angle region (Fig. 4A). On the other hand, the T195 profile exhibited its deepest minimum in the higher-angle region, while a second low-free-energy basin remained present at lower dihedral angles (Fig. 4B).

**Fig. 4.**
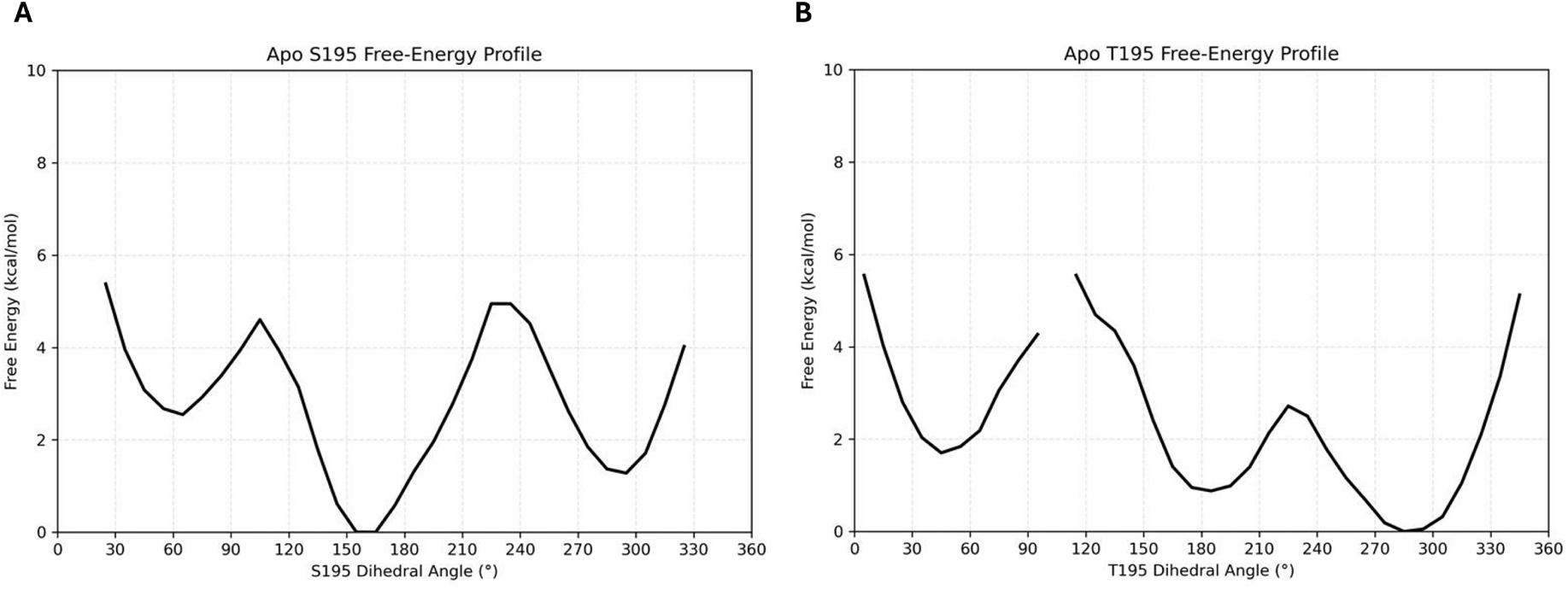
Population-derived free-energy profiles of S195 and T195 from long-timescale apo thrombin simulations. PMFs along the C–C*α*–C*β*–O*γ* dihedral angle are shown for (A) apo S195 and (B) apo T195 thrombin. PMFs were calculated from the combined dihedral-angle distributions of five independent 30 *µ*s unbiased molecular dynamics simulations, corresponding to 150 *µ*s of aggregate sampling per system. Each profile was referenced to its respective free-energy minimum. Bins with zero observed population were masked and are shown as gaps in the free-energy profiles.

To directly assess the agreement between the two simulation approaches, the population-derived free-energy profiles from long-timescale unbiased MD simulations were compared with those obtained from well-tempered metadynamics (Fig. 5). Overall, the two methods exhibited good qualitative agreement in the locations of the major conformational basins for both S195 and T195. For S195, both approaches identified a prominent low-free-energy region near 160-190*^◦^* and a secondary basin near 280-300*^◦^* (Fig. 5A). Similarly, for T195, both methods identified low-energy regions near 180-200*^◦^* and 270-300*^◦^* (Fig. 5B).

**Fig. 5.**
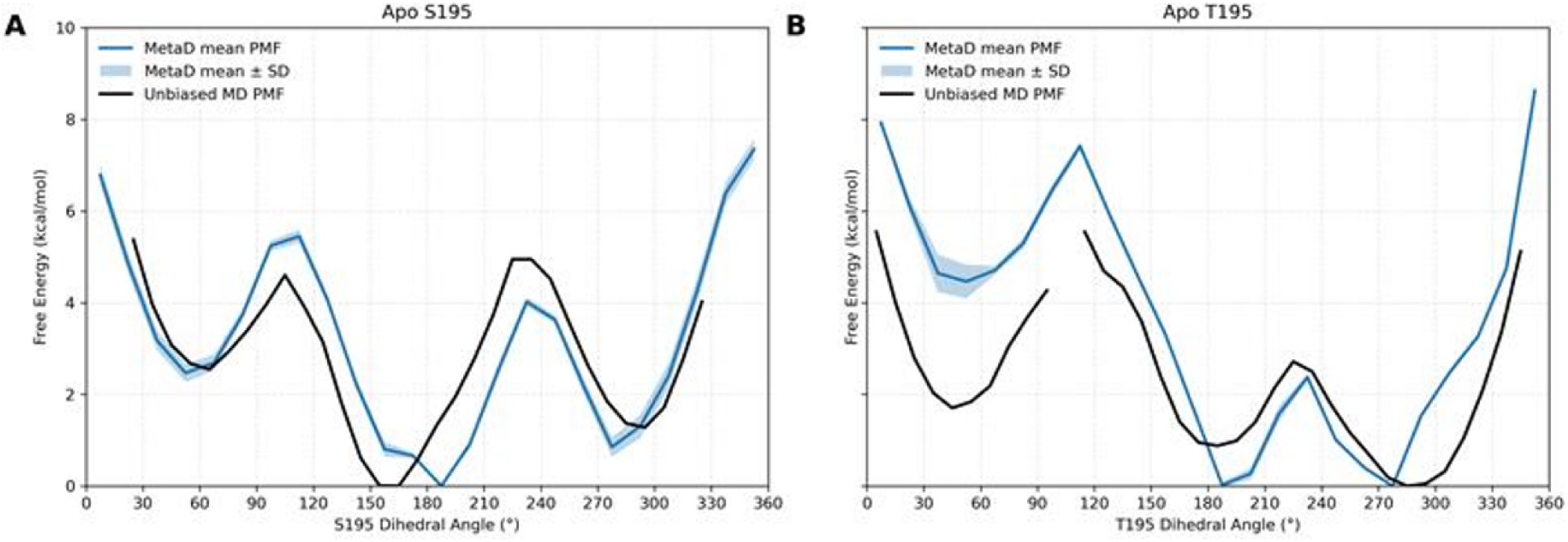
Comparison of free-energy profiles from metadynamics and long-timescale unbiased molecular dynamics simulations of apo S195 and T195 thrombin. PMFs along the C–C*α*–C*β*–O*γ* dihedral angle are shown for (A) apo S195 and (B) apo T195 thrombin. Blue curves represent the mean PMFs obtained from well-tempered metadynamics simulations, with shaded regions indicating the standard deviation over the final 20 ns of the 100 ns simulations. Black curves represent the population-derived PMFs calculated from five independent 30 *µ*s unbiased MD simulations, corresponding to 150 *µ*s of aggregate sampling per system. Each profile was referenced to its respective free-energy minimum. Bins with zero observed population in the unbiased MD profiles were masked and are shown as gaps.

Quantitative comparison of the principal conformational basins further supported the qualitative agreement between the two simulation approaches (Table 1). For S195, the lower-angle free-energy minimum occurred at 187.5*^◦^* in metadynamics and at 165*^◦^* in the unbiased MD simulations, while the higher-angle minima were located at 277.5*^◦^* and at 295*^◦^*, respectively. Both approaches favored the lower-angle basin, with relative free-energy differences of 0.85 and 1.28 kcal/mol for metadynamics and unbiased MD simulations, respectively. For T195, closer agreement was observed in basin positions, with lower-angle minima at 187.5*^◦^* and at 185*^◦^* and higher-angle minima at 277.5*^◦^* and at 285*^◦^* for metadynamics and unbiased MD simulations, respectively. Although metadynamics predicted nearly equivalent free energies for the two T195 basins, unbiased MD simulations favored the higher-angle conformation by approximately 0.88 kcal/mol. The free-energy barriers separating the principal basins also differed quantitatively between the methods. For S195, the barrier from the lower-angle basin to the higher-angle basin was 4.02 kcal/mol in metadynamics and 4.95 kcal/mol in unbiased MD simulations, whereas the corresponding barriers for T195 were 2.35 and 1.84 kcal/mol, respectively. Thus, both approaches consistently predicted a higher barrier for S195 than for T195 along the selected angular pathway, despite differences in the absolute barrier heights.

**Table 1:** Comparison of the principal conformational basin positions, relative free energies, and barrier heights obtained from metadynamics and long-timescale unbiased MD simulations of apo S195 and T195 thrombin. Δ*G* represents the free-energy difference between the higher- and lower-angle basin minima (*G*_higher_ *−G*_lower_). Forward and reverse barriers were calculated relative to the lower- and higher-angle basin minima, respectively, using the intervening free-energy maximum.

| System | Method | Basin position (°) | | $\Delta G$<br>(kcal/mol) | Barrier (kcal/mol) | |
| --- | --- | --- | --- | --- | --- | --- |
|  |  | Lower | Higher |  | Forward | Reverse |
| S195 | MetaD | 187.5 | 277.5 | 0.851 | 4.019 | 3.168 |
| S195 | Unbiased MD | 165.0 | 295.0 | 1.279 | 4.947 | 3.667 |
| T195 | MetaD | 187.5 | 277.5 | -0.024 | 2.347 | 2.371 |
| T195 | Unbiased MD | 185.0 | 285.0 | -0.878 | 1.838 | 2.716 |

Together, the metadynamics and long-timescale unbiased simulations demonstrate that both S195 and T195 can access lower- and higher-angle conformational states in apo thrombin, but the S195T substitution alters their relative conformational preferences. Despite quantitative differences in basin positions, relative free energies, and barrier heights, both approaches independently identified similar principal conformational basins and supported a shift in preference toward the higher-angle state in T195. The convergence of these two distinct sampling approaches strengthens the conclusion that the additional methyl group of T195 modifies the conformational preferences of the catalytic side chain, even in the absence of a ligand.

### Rotameric Preferences Are Associated with Differences in Catalytic-Triad Geometry

The distinct rotameric preferences observed for S195 and T195 raise the question of whether these conformational differences influence the positioning of the catalytic hydroxyl relative to His57. The hydrogen-bond interaction between the hydroxyl group of residue 195 and His57 is an important component of the catalytic triad in serine proteases. To investigate whether the S195T substitution alters this interaction, we examined the distance between His57 N*ɛ*2 and the hydroxyl oxygen of residue 195 across the five independent apo simulations (Fig. **??**). To determine whether differences in this interaction were associated with the distinct rotameric populations of S195 and T195, the analysis was performed separately for the lower-angle (150-210*^◦^*) and higher-angle (260-310*^◦^*) rotameric regions (Fig. 6).

**Fig. 6.**
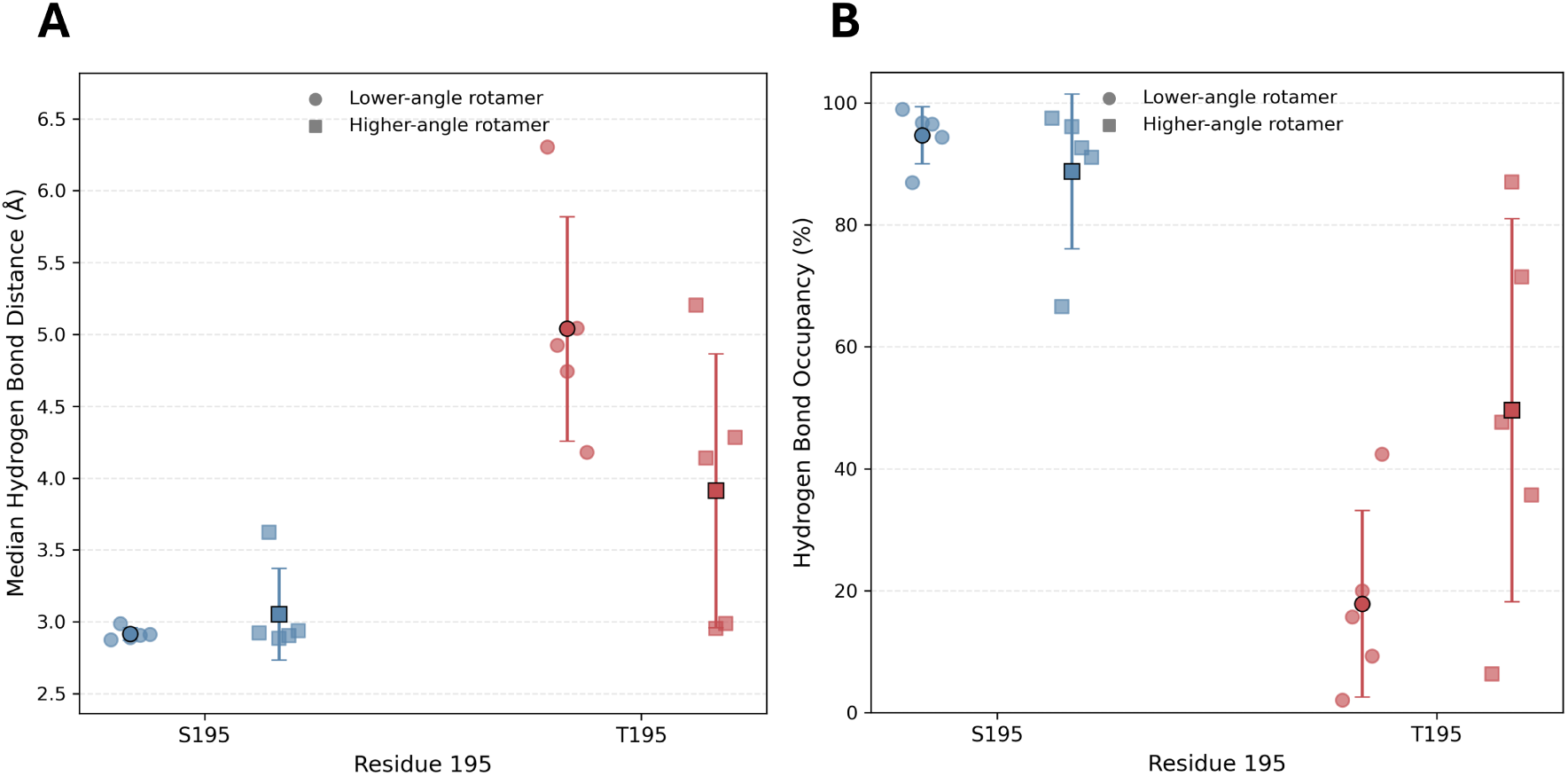
Rotamer-conditioned His57-Ser/Thr195 hydrogen bond distances and distance-based occupancies in apo S195 and T195 thrombin. (A) Median distances between His57 N*ɛ*2 and the hydroxyl oxygen of residue 195 (S195 OG or T195 OG1) and (B) distance-based occupancies calculated using a 4 Å cutoff. Analyses were performed separately for the lower-angle (150-210*^◦^*) and higher-angle (260-310*^◦^*) rotameric regions, defined by the C–C*α*–C*β*-O*γ* dihedral angle of residue 195. S195 and T195 are represented in blue and red, respectively. Circles indicate the lower-angle rotamer, while squares indicate the higher-angle rotamer. Individual points represent values from each of the five independent 30 *µ*s unbiased MD simulations, with larger outlined markers indicating the mean across replicates and error bars representing the standard deviation. Frames outside the defined rotameric regions were excluded from the conditioned analysis.

S195 maintained a relatively short hydrogen bond minimum distance in both rotameric regions. The mean replicate-level median distances were 2.92 *±* 0.04 Å and 3.05 *±* 0.32 Å for the lower- and higher-angle rotamers, respectively (Fig. 6A). Similarly, the percentage of frames with a His57-O195 distance within 4 Å remained high in both regions, averaging 94.71 *±* 4.66% and 88.77 *±* 12.67%, respectively (Fig. 6B). These results indicate that S195 maintains close proximity to His57 across both rotameric regions, despite differences in their overall populations.

In contrast, T195 exhibited greater variability in the His57-O195 distance and reduced distance-based contact occupancy. Within the lower-angle rotamer, the mean replicate-level median distance increased to 5.04 *±* 0.78 Å, with a corresponding occupancy of 17.89 *±* 15.30%. Interestingly, the higher-angle T195 rotamer exhibited a shorter mean replicate-level median distance of 3.91 *±* 0.95 Å and a higher contact occupancy of 49.64 *±* 31.39%. Although this indicates that the higher-angle T195 conformation more frequently positions the hydroxyl near His57 than the lower-angle conformation, its contact occupancy remained considerably lower than that observed for S195 in either rotameric region. The substantial variation among T195 replicates further suggests that the geometry of this interaction is less consistently maintained following the substitution.

Together, these findings indicate that the S195T substitution is associated with changes in the positioning of the catalytic hydroxyl relative to His57 that are not explained solely by differences in rotamer populations. Whereas S195 maintains close His57–hydroxyl proximity across both conformational regions, T195 exhibits less consistent proximity, even within corresponding rotameric states. Although the lower-angle T195 region was sparsely sampled in four of the five replicates, the overall differences suggest that the substitution affects the organization of the catalytic triad. These changes provide a potential structural basis for the preference for serine over threonine at the catalytic nucleophilic position of thrombin.

### The T195 Methyl Group Reorganizes Local Protein Packing

Having established that the S195-to-T195 substitution alters the conformational preferences of the catalytic side chain, we next examined whether the additional methyl group of T195 modifies its local protein environment. Contact occupancies within 5 Å revealed substantial redistribution of interactions with residues nearby position 195 (Fig. 7A). The clearest difference occurred for S214, which contacted the S195 hydroxyl with an occupancy of 88.19 ± 5.06% across the five replicates. In T195, contact of the hydroxyl group (OG1) with S214 decreased to 22.42 ± 22.35%, whereas the T195 methyl group (CG2) contacted S214 with an occupancy of 89.84 ± 8.54%. A similar pattern was observed for V213 in which contact occupancy was 96.38 ± 1.32% for S195 OG, but decreased to 65.74 ± 19.75% for T195 OG1, and remained high for T195 CG2 at 94.93 ± 7.27%. Therefore, for both S214 and V213, introduction of the T195 methyl group was accompanied by reduced contact of the hydroxyl group and increased or maintained contact on the methyl-facing side of the residue.

**Fig. 7.**
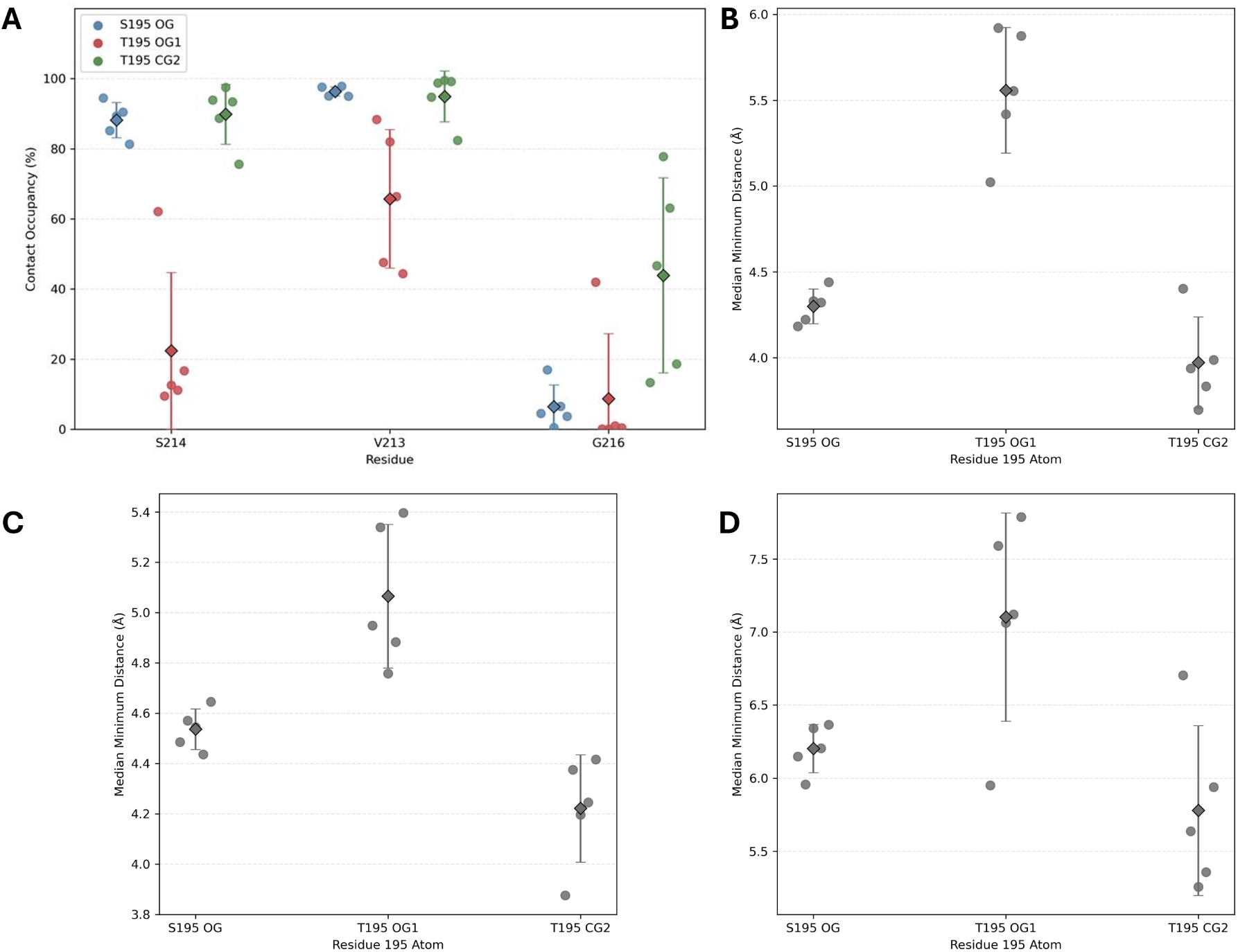
The T195 methyl group reorganizes local packing around residue 195. (A) Contact occupancies of S195 OG (blue), T195 OG1 (red), and T195 CG2 (green) with S214, V213, and G216 during five independent 30 *µ*s apo simulations. Individual points represent replicate occupancies, diamonds indicate the mean across the five replicates, and error bars represent the standard deviation. (B–D) Replicate-level median minimum heavy-atom distances between residue 195 and (B) S214, (C) V213, and (D) G216. Individual points represent the median minimum distance from each replicate, diamonds indicate the mean of the five replicate medians, and error bars represent the standard deviation.

The redistribution was also apparent for G216, although with greater replicate-to-replicate variability. Contact occupancies for S195 OG and T195 OG1 were relatively low, averaging 6.44 ± 6.27% and 8.73 ± 18.59%, respectively, whereas T195 CG2 exhibited a substantially higher mean occupancy of 43.91 ± 27.82% (Fig. 7A). The large variation among T195 replicates indicates that interaction of the methyl group with G216 is more conformationally dependent than the persistent contacts observed with S214 and V213.

Minimum-distance analyses further supported the altered packing environment around T195 (Fig. 7B–D). For S214, the mean of the replicate-level median minimum distances was 4.30 ± 0.10 Å from S195 OG and increased to 5.56 ± 0.37 Å from T195 OG1 (Fig. 7B). In contrast, the corresponding distance from T195 CG2 was 3.97 ± 0.27 Å, with CG2 closer to S214 than OG1 in all five T195 replicates. A similar relationship was observed for V213 (Fig. 7C): the median minimum distance averaged 4.54 ± 0.08 Å for S195 OG, 5.07 ± 0.29 Å for T195 OG1, and 4.22 ± 0.21 Å for T195 CG2. Again, CG2 was closer to V213 than OG1 in each of the five T195 simulations.

The same general tendency extended to G216, although it was less consistent across trajectories (Fig. 7D). Median minimum distances averaged 6.20 ± 0.17 Å for S195 OG, 7.10 ± 0.71 Å for T195 OG1, and 5.78 ± 0.58 Å for T195 CG2. The methyl group was closer to G216 than the T195 hydroxyl in four of the five replicates, consistent with the greater heterogeneity observed in the contact-occupancy analysis. The corresponding minimum-distance trajectories across the five independent long-timescale simulations (Figures **??**, **??**, and **??**) further demonstrate how the introduction of the T195 methyl group alters the dynamic packing environment surrounding the catalytic residue.

To determine whether the differences in local packing between S195 and T195 were solely attributable to their distinct rotameric populations, contact occupancies and minimum distances were further analyzed within the lower-angle (150-210*^◦^*) and higher-angle (260-310*^◦^*) conformational regions (Fig. 8). This analysis allowed the local interactions of S195 and T195 to be compared within corresponding rotameric states, independently of differences in their overall conformational populations. The resulting distributions revealed that differences in local contacts persisted within the same rotameric regions, although the magnitude and nature of these differences depended on the neighboring residue.

**Fig. 8.**
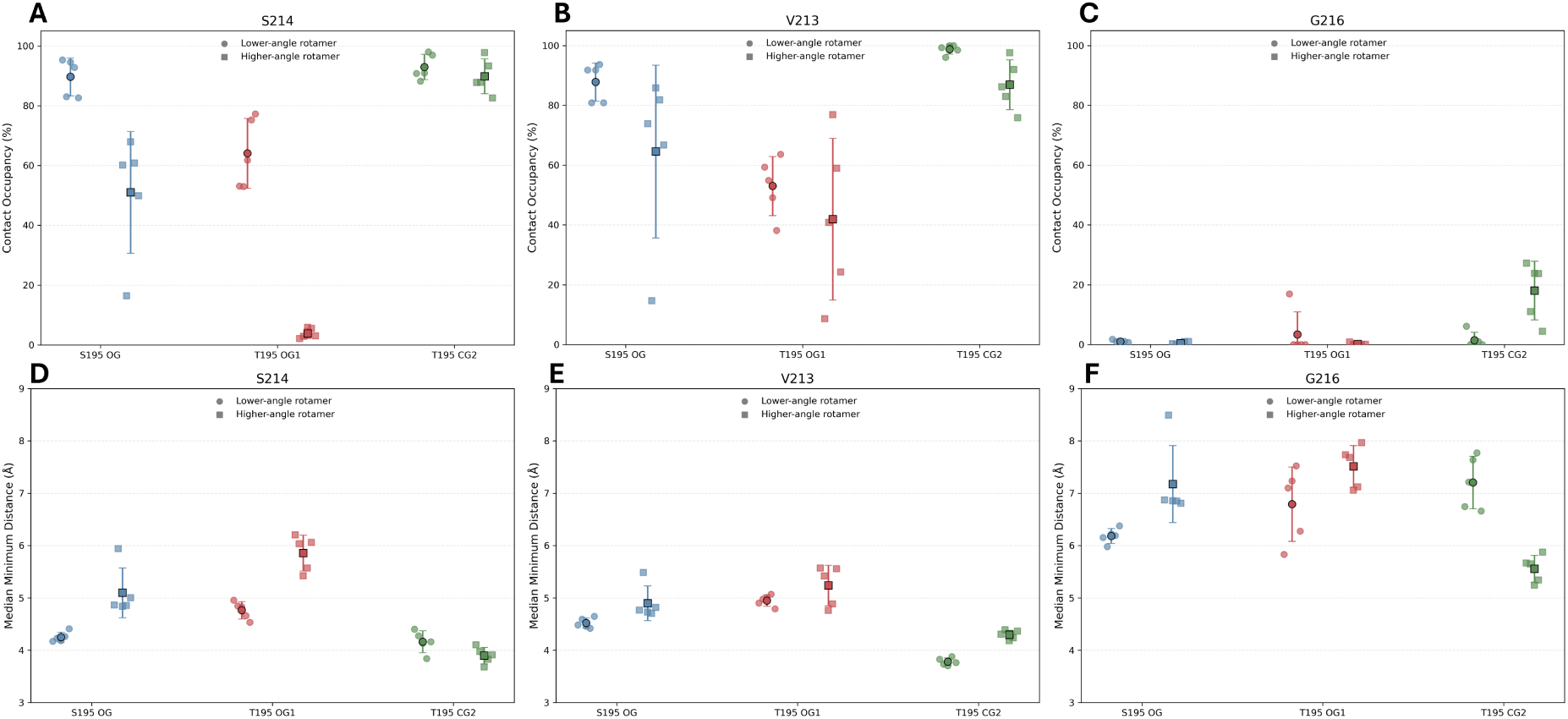
Rotamer-conditioned contact occupancies and minimum distances between residue 195 and neighboring residues in apo S195 and T195 thrombin. Contact occupancies within 5 Å are shown for (A) S214, (B) V213, and (C) G216, while the corresponding median minimum heavy-atom distances are shown for (D) S214, (E) V213, and (F) G216. Analyses were performed separately for the lower-angle (150–210*^◦^*) and higher-angle (260–310*^◦^*) rotameric states, defined using the C–C*α*–C*β*–O*γ* dihedral angle of residue 195. S195 OG, T195 OG1, and T195 CG2 are represented in blue, red, and green, respectively. Circles represent the lower-angle rotamer, while squares represent the higher-angle rotamer. Individual points represent values obtained from each of the five independent 30 *µ*s unbiased MD simulations with larger outlined markers indicating the mean across replicates and error bars representing the standard deviation. Frames outside the defined rotameric regions were excluded from the analyses.

Rotamer-conditioned analysis revealed distinct interaction patterns for S214 and V213 (Fig. 8A,B). Within the lower-angle rotamer, S195 OG maintained higher contact occupancies with both residues than T195 OG1, indicating that the reduced hydroxyl interactions in T195 were not solely explained by the redistribution of rotameric populations. The corresponding minimum-distance analyses supported these differences, with T195 OG1 positioned farther from S214 and V213 than S195 OG within the same conformational region (Fig. 8D,E). In contrast, T195 CG2 maintained high contact occupancies and relatively short distances to both residues in the lower- and higher-angle rotamers, suggesting that the additional methyl group remains closely associated with these neighboring residues across the major conformational states. A different pattern was observed for G216, where T195 CG2 exhibited greater contact occupancy and shorter distances in the higher-angle rotamer than in the lower-angle rotamer (Fig. 8C,F). These observations suggest that the additional methyl group is associated with changes in local packing that extend beyond differences in rotamer populations, while some interactions remain strongly dependent on the side-chain conformation. Although these trends were observed across the rotamer-conditioned analyses, the lower-angle T195 state was sparsely sampled in four of the five independent trajectories, and the corresponding interaction estimates should therefore be interpreted with caution.

Together, the overall and rotamer-conditioned analyses indicate that the S195T substitution is associated with a redistribution of local packing interactions around the catalytic side chain. The persistence of reduced hydroxyl contacts and closer methyl-group interactions with S214 and V213 within corresponding rotameric states suggests that these structural differences are not solely attributable to changes in rotamer populations. Furthermore, the rotamer-dependent interaction of T195 CG2 with G216 highlights how the orientation of the additional methyl group influences its local packing environment. These findings provide a structural context for the altered conformational preferences of S195 and T195 observed in the long-timescale unbiased and metadynamics simulations. Structural representations of the apo S195 and T195 active sites are shown in Figure **??**.

### The Ser195-to-Thr Substitution Alters Side-Chain Solvent Accessibility and Hydration

Since the additional methyl group of T195 altered the local packing environment surrounding residue 195, we next examined whether these structural changes were accompanied by differences in solvent exposure and hydration of the catalytic side chain. Atom-specific solvent-accessible surface area (SASA) was calculated separately for the S195 hydroxyl oxygen (OG), the T195 hydroxyl oxygen (OG1), and the additional T195 methyl carbon (CG2) across the five independent apo simulations (Fig. 9A).

**Fig. 9.**
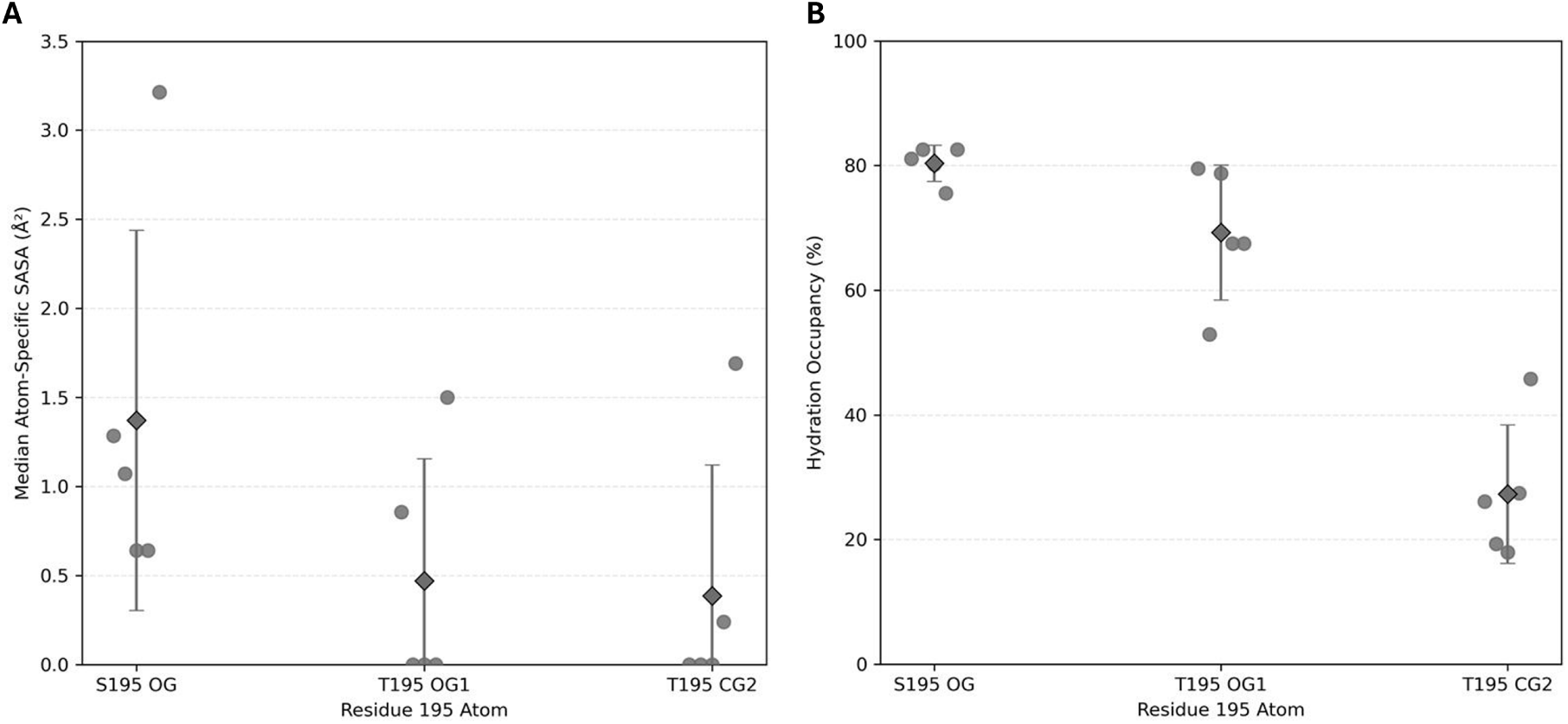
The S195-to-T195 substitution alters side-chain solvent accessibility and water contact. (A) Atom-specific SASA of S195 OG, T195 OG1, and T195 CG2 across five independent 30 *µ*s apo simulations. Individual points represent replicate-level median SASA values, diamonds indicate the mean across replicates, and error bars represent the standard deviation. (B) Water-contact occupancy for S195 OG, T195 OG1, and T195 CG2, defined as the percentage of frames containing at least one water oxygen within 3.5 Å of the indicated atom. Individual points represent replicate occupancies, diamonds indicate the mean across replicates, and error bars represent the standard deviation.

The S195 hydroxyl exhibited greater and more consistent solvent accessibility than the corresponding T195 hydroxyl. The mean of the replicate-level median SASA values was 1.37 ± 1.07 Å^2^ for S195 OG, compared with 0.47 ± 0.68 Å^2^ for T195 OG1. In three of the five T195 replicates, the median OG1 SASA was 0 Å^2^, indicating that the hydroxyl was solvent inaccessible for at least half of the sampled configurations in those trajectories. The T195 methyl group was similarly poorly exposed, with a mean replicate-level median SASA of 0.39 ± 0.74 Å^2^ and median values of 0 Å^2^ in three of the five replicates. These results indicate that both the hydroxyl and methyl groups of T195 experience low and heterogeneous solvent accessibility, consistent with the methyl-side packing observed with neighboring protein residues in Figure 7. The corresponding SASA trajectories across the five independent long-timescale simulations are provided in Figure **??** and illustrate the dynamic changes in solvent accessibility of the catalytic side chain associated with the S195T substitution.

Differences in solvent accessibility were accompanied by changes in local water contact (Fig. 9B). S195 OG maintained at least one water oxygen within 3.5 Å in 80.38 ± 2.89% of trajectory frames, with similar occupancies across all five replicates. Water-contact occupancy for T195 OG1 was lower and substantially more variable, averaging 69.24 ± 10.82%. Thus, although the T195 hydroxyl remained hydrated for a considerable fraction of the simulations, its interaction with the surrounding solvent was less persistent and more dependent on the conformational state sampled by an individual trajectory. Water proximity to the T195 methyl group was substantially lower, with CG2 exhibiting an occupancy of only 27.34 ± 11.11%, consistent with its preferential association with the local protein environment rather than solvent.

To determine whether the differences in solvent accessibility and hydration between S195 and T195 were primarily associated with their distinct rotameric populations, SASA and water-contact occupancies were further analyzed separately within the lower-angle (150-210*^◦^*) and higher-angle (260-310*^◦^*) conformational regions (Fig. 10). This analysis revealed that hydroxyl solvent accessibility varied with rotameric conformation and that the differences between S195 and T195 were not uniform across the two regions. For S195 OG, the mean replicate-level median SASA increased from 1.33 *±* 1.11 Å^2^ in the lower-angle region to 2.10 *±* 1.11 Å^2^ in the higher-angle region (Fig. 10A). In contrast, T195 OG1 exhibited a higher mean replicate-level median SASA in the lower-angle region (3.11 *±* 1.96 Å^2^) than in the higher-angle region (1.11 *±* 1.64 Å^2^). The T195 methyl group remained poorly solvent accessible in both conformational regions, with mean replicate-level median SASA values of 0 and 0.63 *±* 0.78 Å^2^ for the lower- and higher-angle rotamers, respectively. These results suggest that the relationship between the S195T substitution and hydroxyl solvent exposure is strongly dependent on side-chain conformation.

**Fig. 10.**
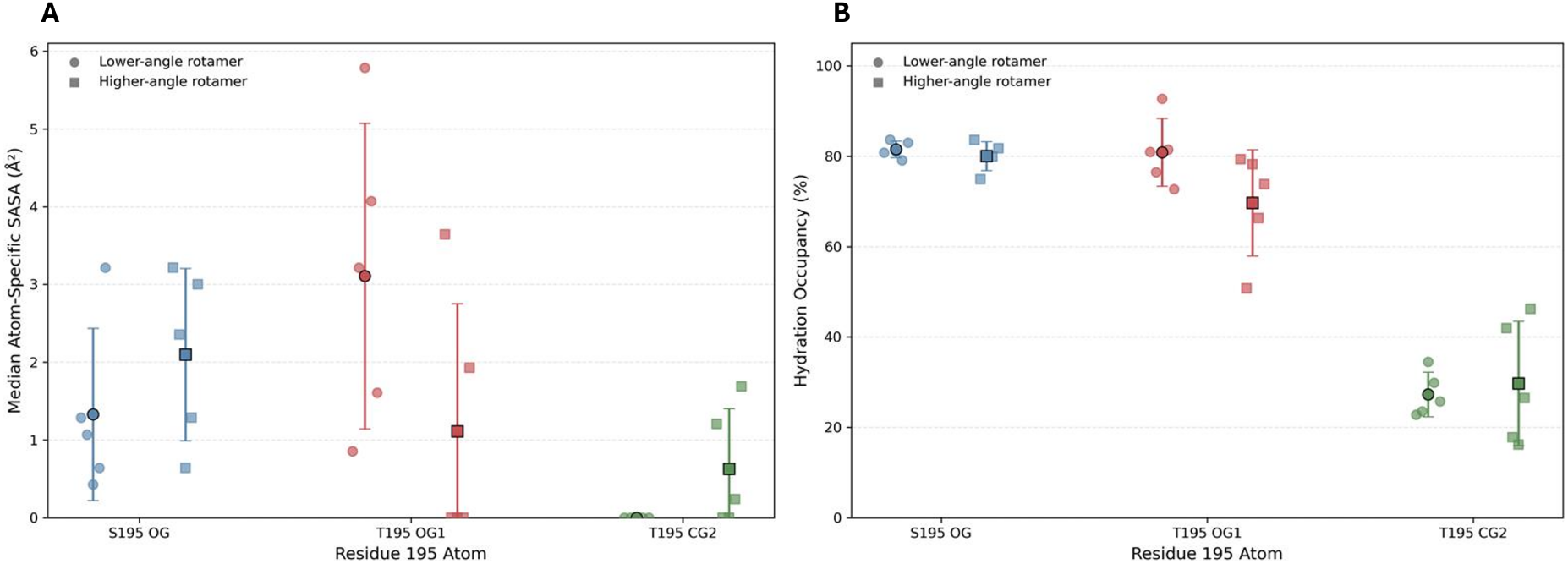
Rotamer-conditioned solvent accessibility and water-contact occupancy of residue 195 in apo S195 and T195 thrombin. (A) Atom-specific SASA and (B) water-contact occupancy of S195 OG, T195 OG1, and T195 CG2, calculated separately for the lower-angle (150-210*^◦^*) and higher-angle (260-310*^◦^*) rotameric regions. Rotameric states were defined using the C–C*α*–C*β*–O*γ* dihedral angle of residue 195. Water-contact occupancy was defined as the percentage of frames containing at least one water oxygen within 3.5 Å of the indicated atom. S195 OG, T195 OG1, and T195 CG2 are shown in blue, red, and green, respectively. Circles represent the lower-angle rotamer, while squares represent the higher-angle rotamer. Individual points represent replicate-level median SASA values (A) or water-contact occupancies (B) from five independent 30 *µ*s unbiased MD simulations. Larger outlined markers indicate the mean across replicates, and error bars represent the standard deviation. Frames outside the defined rotameric regions were excluded from these analyses.

Rotamer-conditioned water-contact analysis further demonstrated that the hydration behavior of the catalytic hydroxyl depends on side-chain conformation (Fig. 10B). S195 OG maintained similar water-contact occupancies in the lower- and higher-angle regions, averaging 81.53 *±* 1.84% and 80.03 *±* 3.22%, respectively. T195 OG1 exhibited comparable hydration to S195 OG within the lower-angle region, with an occupancy of 80.86 *±* 7.53%, but showed reduced water contact in the higher-angle region, averaging 69.68 *±* 11.76%. In contrast, T195 CG2 exhibited relatively low water-contact occupancies in both conformational regions, averaging 27.31 *±* 4.90% and 29.72 *±* 13.75% for the lower- and higher-angle rotamers, respectively. These findings indicate that the reduced overall hydration of the T195 hydroxyl is associated, at least in part, with its greater sampling of the higher-angle conformation, where its water-contact occupancy is lower than that of S195 OG. However, the lower-angle T195 rotamer was sparsely sampled in four of the five independent trajectories, with only 51-97 frames assigned to this region in each of those replicates. Therefore, the corresponding rotamer-conditioned SASA and hydration estimates should be interpreted cautiously.

Together, the overall and rotamer-conditioned SASA and water-contact analyses demonstrate that the S195T substitution is associated with changes in the local solvation environment of the catalytic side chain. Although the overall distributions indicate reduced solvent accessibility and hydration of the T195 hydroxyl compared with S195, the rotamer-conditioned analyses reveal that these differences vary across conformational states. In particular, hydroxyl water-contact occupancies were comparable within the lower-angle region, whereas T195 exhibited reduced hydration in the higher-angle region. The additional T195 methyl group remained largely solvent inaccessible and exhibited relatively low water-contact occupancy in both rotamers, consistent with its preferential packing against neighboring protein residues. These findings suggest that the altered solvation environment of T195 reflects both differences in conformational populations and rotamer-dependent interactions with the surrounding protein and solvent, providing further structural context for the conformational preferences observed in the enhanced-sampling and long-timescale unbiased simulations.

## Conclusion

In this study, enhanced-sampling and microsecond-long MD simulations were used to characterize how substituting the catalytic S195 residue with T195 alters the conformational energetics and the local molecular environment of thrombin. Well-tempered metadynamics revealed distinct free-energy landscapes for S195 and T195 in both the apo and PPACK-bound states. In the apo enzyme, S195 preferentially sampled the lower-angle rotameric state, whereas the corresponding T195 populated two major low-energy regions with nearly equivalent energetic stability. PPACK binding shifted both residues toward the lower-angle region, with substantially stronger energetic discrimination against the higher-angle state for S195. Long-timescale unbiased simulations further demonstrated that S195 predominantly sampled the lower-angle conformational region, whereas T195 exhibited greater replicate-to-replicate heterogeneity and increased sampling of the higher-angle state. Overall, our results are consistent with the findings reported by Pelc et al. in which apo S195 tends to sample lower-angle dihedral states, apo T195 tends to sample higher-angle dihedral states, and the binding of PPACK in either condition causes a shift in the dihedral angles to resemble that of apo S195.^12^

The differences in conformational preferences between S195 and T195 were also accompanied by changes in the geometry of the catalytic triad. Analysis of the hydrogen bond between His57 N*ɛ*2 and the hydroxyl oxygen of residue 195 revealed that S195 maintained relatively short distances and high distance-based occupancies across both rotameric regions. In contrast, T195 exhibited longer and more variable distances, with substantially reduced occupancies in both regions. Interestingly, the higher-angle T195 rotamer maintained closer proximity to His57 than the lower-angle rotamer, although its contact occupancy remained lower than that observed for S195. The reduced consistency of His57–hydroxyl proximity in T195 provides a plausible structural connection for the preference for serine at the catalytic position of thrombin.

Analysis of the local environment surrounding residue 195 showed that the additional T195 methyl group redistributes protein packing and solvation rather than producing a simple static steric clash. Contacts with neighboring residues were shifted toward the T195 methyl group, while the T195 hydroxyl was positioned farther from several nearby residues. Overall, the T195 hydroxyl displayed reduced and more heterogeneous solvent accessibility and water contact relative to S195, while the methyl group remained largely buried and comparatively poorly hydrated. However, rotamer-conditioned analyses revealed that these differences depend on side-chain conformation, with hydroxyl hydration being comparable between S195 and T195 in the lower-angle rotamer but reduced for T195 in the higher-angle rotamer. Together, these findings suggest that the S195T substitution alters the conformational free-energy landscape in association with changes in local packing and solvation. Combined with the differences in catalytic-triad geometry, these changes provide a dynamic molecular framework for understanding the strong preference for serine over threonine at the catalytic nucleophilic position of thrombin.

## Acknowledgement

The research reported in this work was supported by the National Institute of General Medical Sciences of the National Institutes of Health under award number R35GM147423, as well as by the National Science Foundation grant CHE 1945465 and the Arkansas Biosciences Institute. This research is part of the Frontera computing project by LRAC at the Texas Advanced Computing Center (TACC) through Grant CHE21003, made possible by the National Science Foundation award OAC-1818253. Anton 3 computer time was provided by the Pittsburgh Supercomputing Center (PSC) through Grant 1R24GM154042 from the National Institutes of Health. The Anton 3 machine at PSC is made available by D. E. Shaw Research. This work is also supported by the Arkansas High-Performance Computing Center which is funded through multiple National Science Foundation grants and the Arkansas Economic Development Commission.

