## Supplemental Information for "Free-Energy and Molecular Dynamics Analysis of Ser195 and Thr195 Rotameric States in Thrombin"

Before obtaining the final free-energy profiles from the well-tempered metadynamics simulations, convergence of the biased simulations was assessed by examining the extent to which the PMF changed throughout the final 20 ns of each 100 ns trajectory. Free-energy profiles were extracted at 2 ns intervals from 82 to 100 ns for the apo S195, apo T195, PPACK-bound S195, and PPACK-bound T195 systems. As shown in Figure S1, the PMF profiles remained consistent throughout this interval, indicating that the conformational free-energy landscapes were stable during the final sampling period. These results support the use of the final 20 ns of each metadynamics trajectory for the averaged PMF calculations reported in the main manuscript.

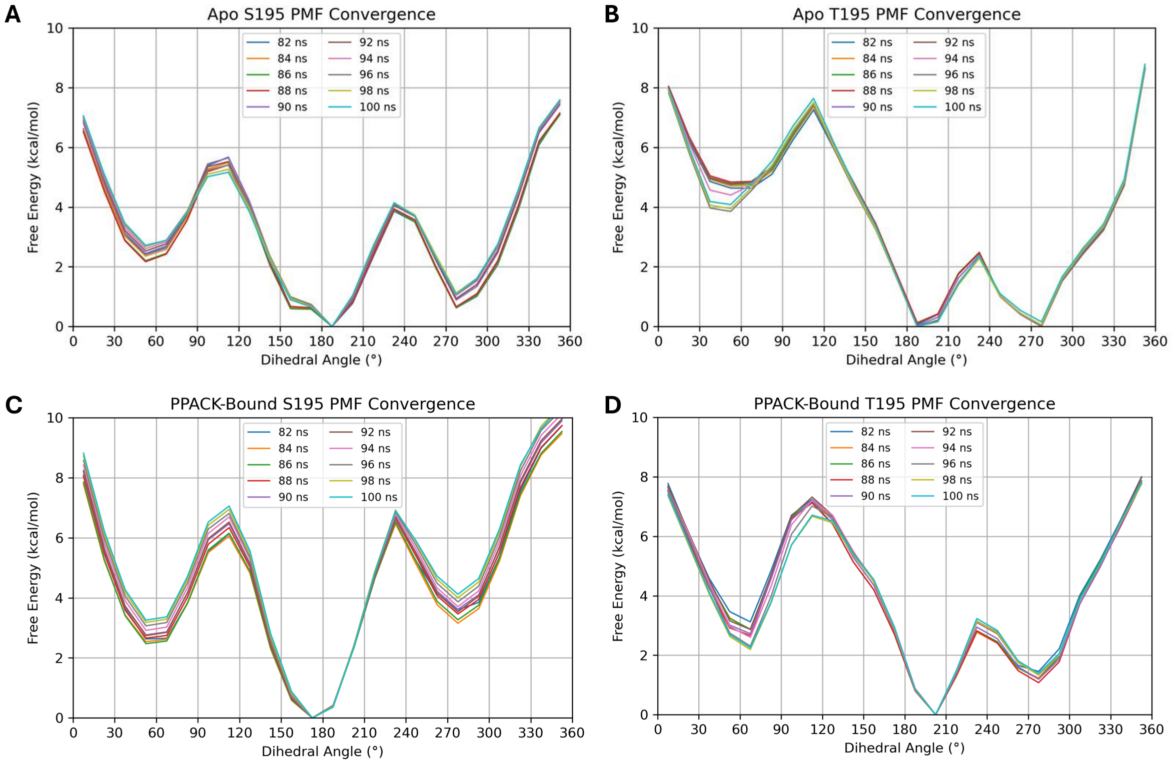


**Fig. S1.** Convergence of well-tempered metadynamics free-energy profiles. Free-energy profiles obtained at 2 ns intervals during the final 20 ns (80–100 ns) of the 100 ns well-tempered metadynamics simulations are shown for (A) apo S195, (B) apo T195, (C) PPACK-bound S195, and (D) PPACK-bound T195 thrombin.

In addition to comparing PMF profiles obtained during the final 20 ns of the simulations, convergence was further evaluated by monitoring the change in relative free energies over time at selected dihedral angles corresponding to regions with greater variation in the free-energy landscape. These regions were selected because they exhibited slower convergence during the well-tempered metadynamics simulations. As shown in Figure S2, the relative free energies at these positions continued to fluctuate during the latter stages of sampling; however, the fluctuations did not result in systematic changes to the overall free-energy landscape.

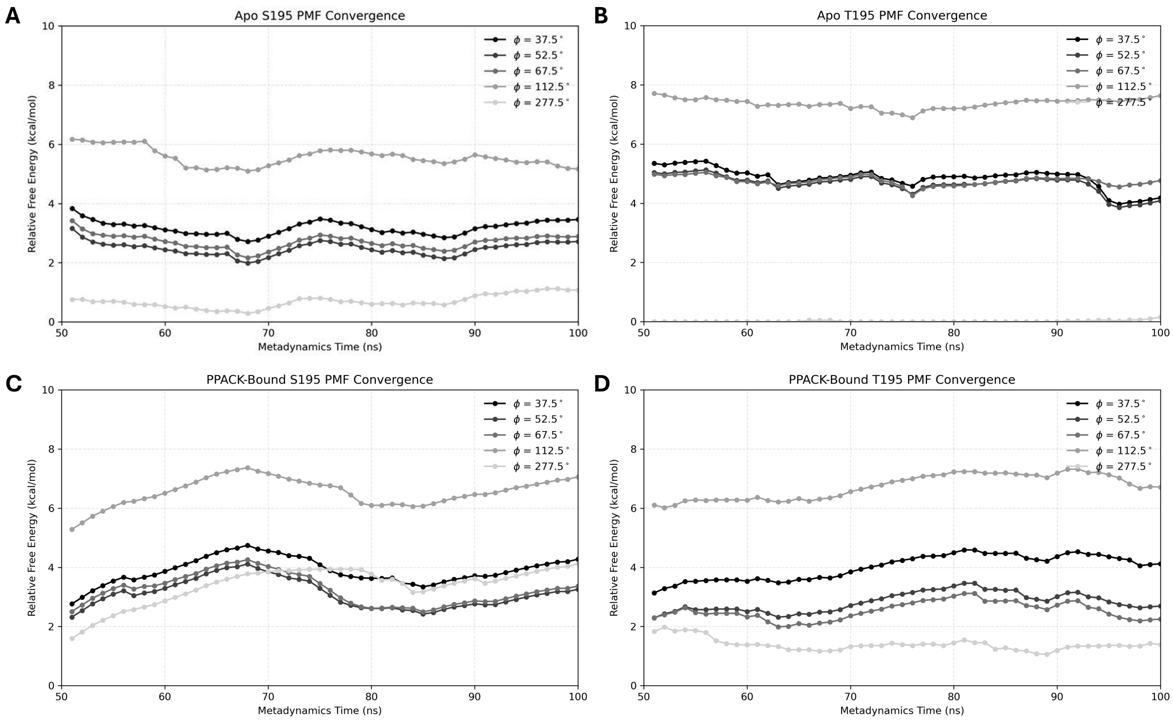


**Fig. S2.** Time evolution of selected relative free-energy values for slowly converging dihedral angle points during well-tempered metadynamics simulations. Relative free energies were monitored at five representative dihedral angles (37.5°,52.5°,67.5°,112.5°, and 277.5°) throughout the well-tempered metadynamics simulations for (A) apo S195, (B) apo T195, (C) PPACK-bound S195, and (D) PPACK-bound T195 thrombin.

Together with the overlap of the PMF profiles obtained during the final 20 ns of sampling (Fig. S1), these analyses indicate that the simulations sufficiently sampled the relevant conformational states and that additional simulation time would be unlikely to substantially alter the calculated free-energy landscapes.

To visualize the conformational sampling underlying the dihedral angle distributions reported in the main manuscript, the time evolution of the catalytic residue side-chain dihedral angle was analyzed for each independent 30 $\mu$s apo thrombin simulation. The trajectories from the five S195 replicates and the five T195 replicates are shown in Figure S3. The individual replicate trajectories demonstrate that S195 predominantly samples a similar conformational region across independent simulations, whereas T195 exhibits greater variability in the sampled dihedral states between replicates. These trajectory-level observations are consistent with the population distributions presented in the main manuscript.

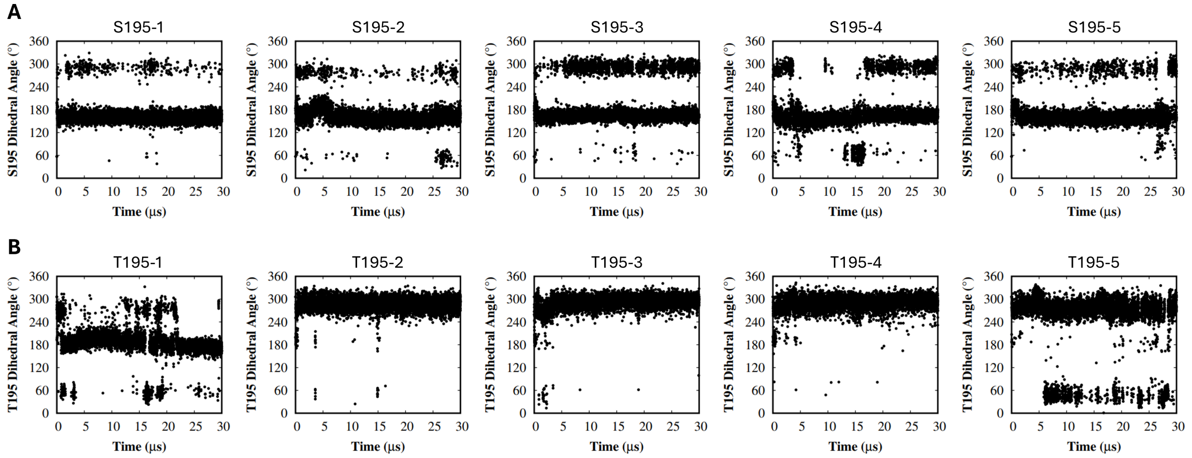


**Fig. S3.** Time evolution of catalytic residue side-chain dihedral angles during long-timescale apo thrombin simulations. The dihedral angle trajectories from five independent 30 $\mu$s simulations are shown for (A) S195 and (B) T195. Individual replicates are arranged from left to right (replicates 1–5).

To further characterize the interaction between His57 and the catalytic hydroxyl of residue 195, the distance between His57 N$\epsilon$2 and S195 OG or T195 OG1 was examined throughout the five independent apo simulations (Fig. S4). In the S195 simulations, the hydrogen bond distance generally remained close to 3 Å, with occasional fluctuations to larger distances. In contrast, the T195 simulations exhibited greater variability, with several replicates showing prolonged periods of increased separation between His57 and the catalytic hydroxyl. These observations suggest that the S195T substitution is associated with reduced consistency in maintaining close proximity between the two residues. The rotamer-conditioned analysis presented in the main text further characterizes these differences by examining the interaction separately within the lower- and higher-angle conformational regions.

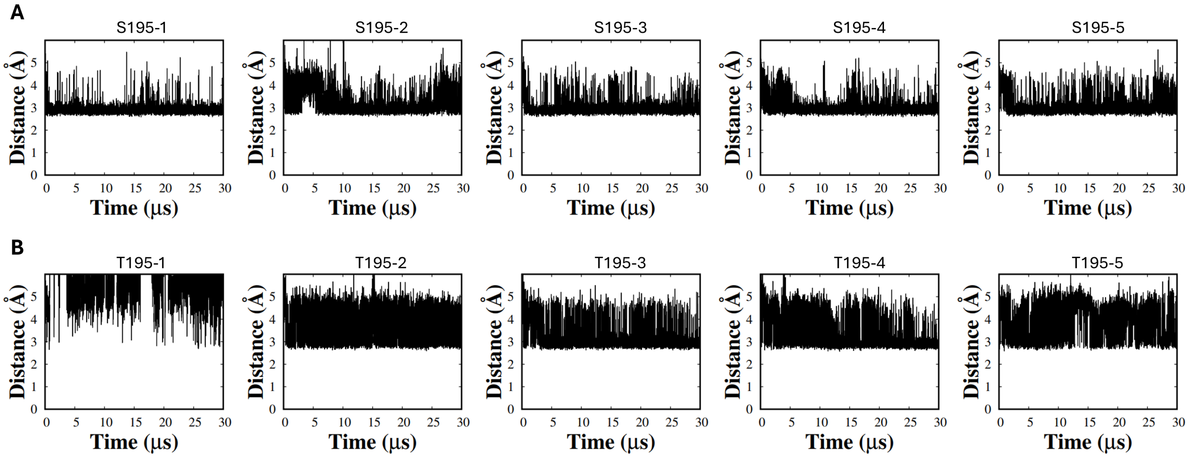


**Fig. S4.** Time evolution of the His57–Ser/Thr195 hydrogen bond distance in apo S195 and T195 thrombin. Distances between His57 N$\epsilon$2 and the hydroxyl oxygen of residue 195 (S195 OG or T195 OG1) are shown for five independent 30 $\mu$s unbiased MD simulations of each system. The upper row shows S195 replicates 1–5, while the lower row shows T195 replicates 1–5.

To further characterize the local structural changes associated with the S195T substitution, minimum distances between residue 195 and neighboring residues were monitored throughout the five independent 30 $\mu$s apo thrombin simulations. The analysis focused on V213, S214, and G216, which are located near the catalytic residue and contribute to the local active-site environment. For each residue, distances were calculated separately for the catalytic hydroxyl atoms of S195 (OG) and T195 (OG1), as well as the additional methyl carbon introduced by the T195 substitution (CG2). The resulting distance trajectories for S214, V213, and G216 are shown in Figures S5, S6, and S7, respectively. These analyses provide a trajectory-level view of the dynamic packing interactions surrounding residue 195 and complement the contact occupancy analysis presented in the main manuscript.

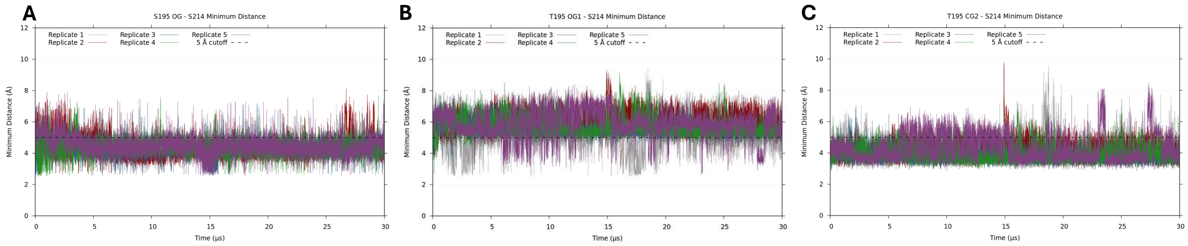


**Fig. S5.** Time evolution of minimum distances between residue 195 and S214 during long-timescale simulations. Minimum distances were calculated between S214 and the catalytic residue atoms S195 OG (A), T195 OG1 (B), and T195 CG2 (C) across five independent 30 $\mu$s apo simulations. The dashed line indicates the 5 Å contact cutoff used for interaction analysis. Individual traces represent independent simulation replicates.

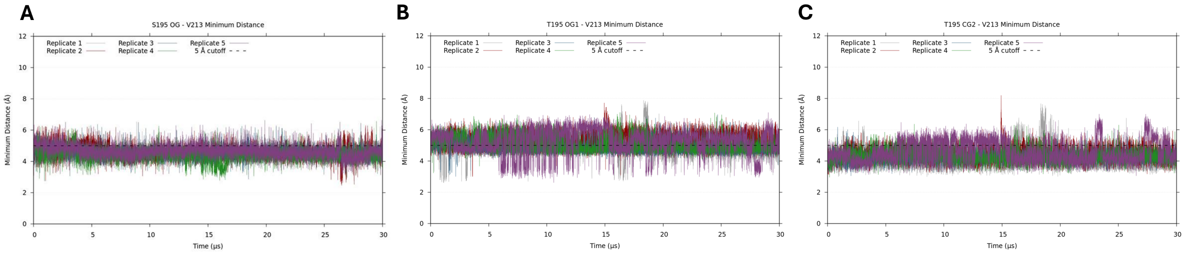


**Fig. S6.** Time evolution of minimum distances between residue 195 and V213 during long-timescale simulations. Minimum distances were calculated between V213 and the catalytic residue atoms S195 OG (A), T195 OG1 (B), and T195 CG2 (C) across five independent 30 $\mu$s apo simulations. The dashed line indicates the 5 Å contact cutoff used for interaction analysis. Individual traces represent independent simulation replicates.

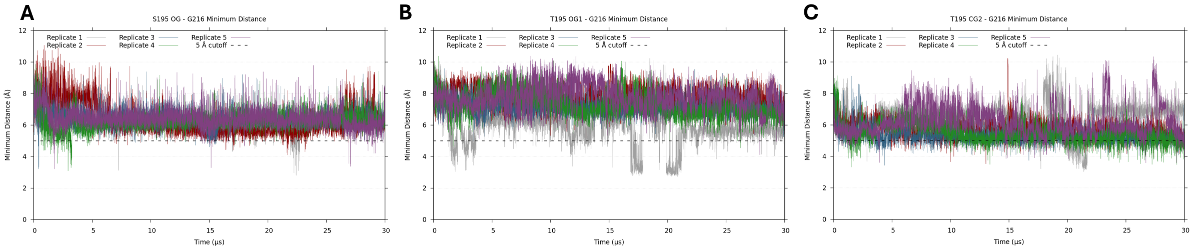


**Fig. S7.** Time evolution of minimum distances between residue 195 and G216 during long-timescale simulations. Minimum distances were calculated between G216 and the catalytic residue atoms S195 OG (A), T195 OG1 (B), and T195 CG2 (C) across five independent 30 $\mu$s apo simulations. The dashed line indicates the 5 Å contact cutoff used for interaction analysis. Individual traces represent independent simulation replicates.

To provide a structural view of the local environment surrounding the catalytic residue, crystal structures of apo S195 and apo T195 thrombin were examined. Representations of the apo S195 and apo T195 active sites are shown in Figure S8 which was generated using VMD.[26] The catalytic residue and neighboring residues analyzed in the contact and minimum-distance calculations are highlighted. In the T195 structure, the additional methyl group introduced by the S195T substitution (CG2) is shown to illustrate the additional steric feature present within the catalytic pocket compared with S195.

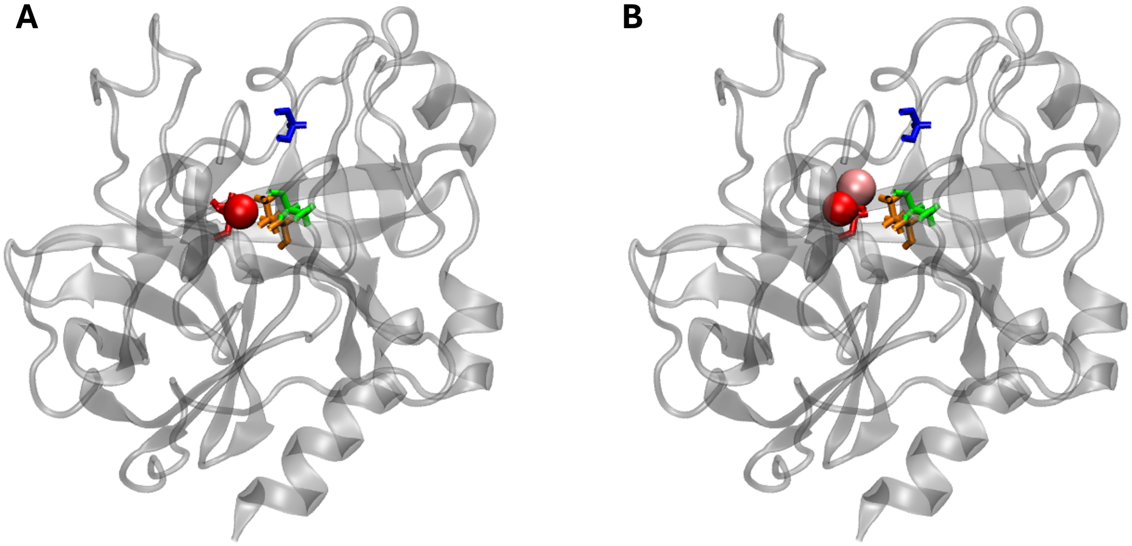


**Fig. S8.** Structural comparison of the apo S195 and apo T195 thrombin active sites. Representative structures of (A) apo S195 thrombin and (B) apo T195 thrombin are shown with the catalytic residue 195 highlighted in red. The catalytic hydroxyl oxygen is shown as a red van der Waals (VDW) sphere for S195 (OG) and T195 (OG1), while the additional methyl carbon introduced by the S195T substitution (CG2) is shown as a pink VDW sphere. Neighboring residues VAL213, SER214, and GLY216 are shown as orange, green, and blue sticks, respectively. The surrounding protein backbone is displayed as a transparent gray NewCartoon representation.

To examine whether the S195T substitution alters the solvent exposure of the catalytic side chain, SASA was monitored throughout the five independent 30 $\mu$s apo thrombin simulations. SASA values were calculated separately for the catalytic hydroxyl atoms of S195 (OG) and T195 (OG1), as well as the additional methyl carbon introduced by the T195 substitution (CG2). The time evolution of these values is shown in Figure S9 and provides a trajectory-level view of changes in local solvent accessibility associated with the mutation.

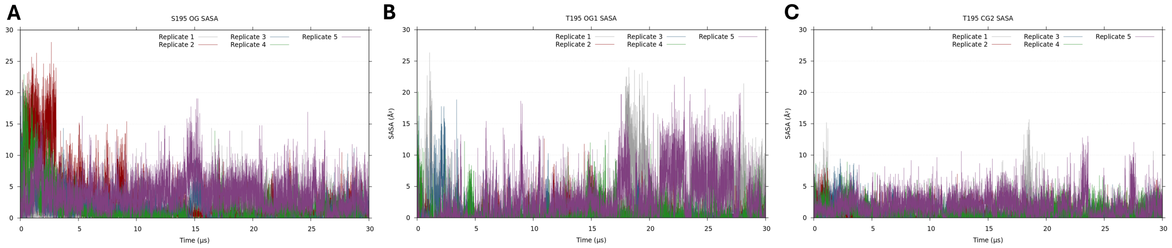


**Fig. S9.** Time evolution of SASA for catalytic residue 195 during long-timescale apo thrombin simulations. SASA values were calculated for (A) the S195 hydroxyl oxygen (OG), (B) the T195 hydroxyl oxygen (OG1), and (C) the additional methyl carbon introduced by the S195T substitution (CG2). The trajectories from five independent 30 $\mu$s simulations are shown for each atom.
